# Burst firing is required for induction of Hebbian LTP at lateral perforant path to hippocampal granule cell synapses

**DOI:** 10.1101/2022.12.06.519388

**Authors:** Yoonsub Kim, Sooyun Kim, Won-Kyung Ho, Suk-Ho Lee

**Author notes:** Corresponding authors *Correspondence* Won-Kyung Ho M.D., Ph.D., Department of Physiology, Seoul National University College of Medicine. 103 Daehak-ro, Jongno-gu, Seoul, 03080, Republic of Korea, e-mail address, Suk-Ho Lee M.D., Ph.D., Department of Physiology, Seoul National University College of Medicine. 103 Daehak-ro, Jongno-gu, Seoul, 03080, Republic of Korea. Competing interests* The authors declare that they have no competing interests.

## Abstract

High frequency burst firing is critical in summation of back-propagating action potentials (APs) in dendrites, which may greatly depolarize dendritic membrane potential. The physiological significance of burst firings of hippocampal dentate GCs in synaptic plasticity remains unknown. We found that GCs with low input resistance could be categorized into regular-spiking (RS) and burst-spiking (BS) cells based on their initial firing frequency (F_init_) upon somatic rheobase current injection, and investigated how two types of GCs differ in long-term potentiation (LTP) induced by high-frequency lateral perforant pathway (LPP) inputs. Induction of Hebbian LTP at LPP synapses required at least three postsynaptic APs at F_init_ higher than 100 Hz, which was met in BS but not in RS cells. The synaptically evoked burst firing was critically dependent on persistent Na^+^ current, which was larger in BS than RS cells. The Ca^2+^ source for Hebbian LTP at LPP synapses was primarily provided by L-type calcium channels. In contrast, Hebbian LTP at medial PP synapses was mediated by T-type calcium channels, and could be induced regardless of cell types or F_init_ of postsynaptic APs. These results suggest that intrinsic firing properties affect synaptically driven firing patterns, and that bursting behavior differentially affects Hebbian LTP mechanisms depending on the synaptic input pathway.

## Introduction

The dentate gyrus (DG) is the first layer at which synaptic inputs from the entorhinal cortex arrive among the hippocampal trisynaptic loop, and has been implicated in pattern separation and conjunctive representations of spatial (context) and non-spatial (items and events) information (Lee & Jung, 2017). The afferent fibers from lateral and medial entorhinal cortex (LEC and MEC) layer II give rise to lateral and medial perforant pathways (LPP and MPP). LPP and MPP carry relatively more non-spatial and spatial information to DG, and innervate distal and intermediate parts of granule cell dendrites in the DG, respectively. Synaptic plasticity-based competitive learning together with the integration of spatial and non-spatial inputs in dentate granule cells (GCs) was recently proposed to underlie the progressive refinement of spatial representation in DG (Kim *et al*., 2020).

Heterogeneity in cellular excitability could be one of key mechanisms underlying recruitment of principal cells to a neuronal ensemble or an engram for representation and formation of memories (Pignatelli *et al*., 2019). Recent in vivo recordings of GCs in DG revealed that a majority of GC spikes occurred in bursts, and that active GCs, which comprised only a minor subset of GCs, were morphologically mature and distinct from silent GCs (Pernía-Andrade & Jonas, 2014; Diamantaki *et al*., 2016; Zhang *et al*., 2020). While these studies imply heterogeneity among dentate GCs, it remains to be understood how the difference in the excitability among heterogeneous groups of mature GCs is related to the difference in synaptic plasticity. Previously, it was noted that the initial frequency (F_init_) of first two action potentials (APs) upon somatic current injection is higher than the rest APs in mature GCs, and T-type voltage dependent Ca^2+^ channels (T-VDCCs) contribute to the burst firing (Dumenieu *et al*., 2018). Burst firing enhances not only the reliability of pre-synaptic glutamate release (Lisman, 1997), but also postsynaptic Ca^2+^ signaling required for synaptic plasticity (Kampa *et al*., 2006; Letzkus *et al*., 2006). Consistently, burst firing of principal cells plays diverse roles in different cortical regions such as place field formation in CA1 (Bittner *et al*., 2017), initiation of sharp waves in CA3 (Hunt *et al*., 2018), and switching thalamic network states for relaying subcortical inputs (Llinás & Steriade, 2006). Moreover, somatic firings above at above a critical frequency has been found to greatly depolarize the dendtritic membrane potential through summation of back-propagating APs in neocortical pyramidal cells (Larkum *et al*., 1999).

In the present study, we found that there are two types of mature GCs displaying low input resistance (< 200 MΩ) in young rats, based on whether the F_init_ of APs elicited by rheobase current injection is higher than 50 Hz or not, referred to as burst-spiking (BS) and regular-spiking (RS) GCs. Similar heterogeneity has been found in CA3 too (Hunt *et al*., 2018). While burst firings of GCs are expected to enhance spike transfer from DG to CA3 due to strong short-term facilitation, it is little understood whether burst firing of GCs has any effect on the input side, that is on the long-term potentiation (LTP) at the LPP or MPP-to-GC synapses. To address this issue, we compared LTP at LPP and MPP synapses between RS and BS cells. We show that Hebbian LTP occurs only at LPP synapses to BS cells but not at those to RS cells, while BS and RS cells do not differ in Hebbian LTP at MPP synapses. The present study on the ionic mechanisms underlying LTP at LPP and MPP synapses to RS and BS-GCs revealed that high and low voltage-activated VDCCs play a key role in LTP induction at LPP and MPP synapses, respectively. These results suggest that activation of L-VDCCs requires high frequency AP firing, providing an insight into why Hebbian LTP at LPP-GC synapses is induced preferentially in BS cells.

## Results

### Characteristics and distribution of two types of mature GCs

Burst firing of dentate GCs has been observed both in vivo (Pernía-Andrade & Jonas, 2014) and ex vivo (Dumenieu *et al*., 2018), but its physiological significance in synaptic plasticity is not well understood. We examined firing patterns of mature GCs that have input resistance (R_in_) less than 200 MΩ in response to somatic current injection. When we applied a step current just above action potential (AP) threshold (rheobase current) for 1 s in whole-cell current clamp mode, a group of cells generated APs in bursts, doublet in majority (82.8%, 18 of 22) and sometimes triplet (18.2%, 4 of 22), while others showed regularly spiking patterns (Fig. 1A). The histogram of initial firing frequency (F_init_) showed bimodal distribution (Fig. 1B), so that we nominated cells with F_init_ under 50 Hz as regular-spiking (RS, gray), while cells with F_init_ over 50 Hz as burst-spiking (BS, red) neurons. The mean value for F_init_ was 10.6 ± 2.2 Hz (n = 18) in RS-GCs and 147.1 ± 11.2 Hz (n = 22) in BS-GCs. As the injection current increased, F_init_ increased in RS-GCs, and the difference of F_init_ between RS- and BS-GCs gradually disappeared (Fig. 1Ca). Despite the remarkable difference in F_init_, the number of APs during 1 s depolarization was not significantly different between two groups (Fig. 1Cb). Analyses of AP shapes revealed that the threshold voltage for AP generation was lower, AP duration was longer, and afterhyperpolarization (AHP) was smaller in BS-GCs compared to those in RS-GCs (Fig. 1D). No significant difference was found in passive electrical properties such as input resistance (R_in_) and resting membrane potential (RMP) (Fig. 1E). Interestingly, in DG-GCs that have R_in_ more than 200 MΩ, which are less mature according to the criteria of maturation (Schmidt-Hieber *et al*., 2004; Kim *et al*., 2018), bursting was very rarely observed (Fig. 1F), suggesting that burst firing is a characteristic feature of fully mature DG-GCs.

**Fig. 1.**
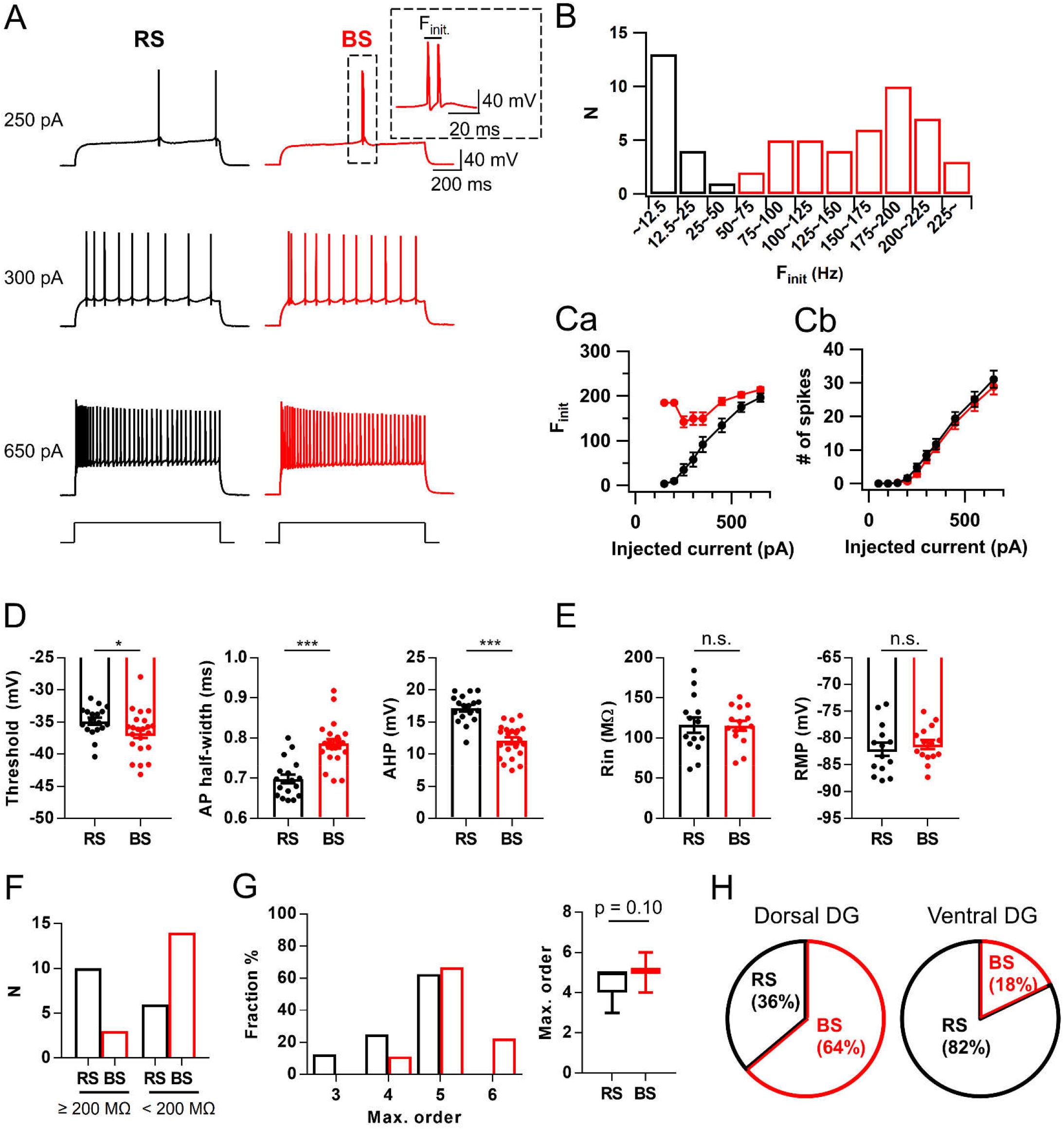
Intrinsic properties of regular-spiking (RS) and burst-spiking (BS) mature granule cells. **A**: Representative voltage responses of RS (black) and BS (red) cells to somatic current injection of 250, 300 and 650 pA (1s duration). *Inset*, initial firing of BS-GCs at expanded time scale. Initial firing frequency (F_init_) was measured as the frequency of first two APs at rheobase current injection. **B**: Bimodal distribution of F_init_ among mature GCs. Mature GCs were divided into RS and BS with the reference frequency of 50 Hz (RS/BS, n = 18/42). **Ca-b**: F_init_ (*a*) and spike numbers (*b*) of RS and BS cells as a function of injected current amplitude (from 50 to 650 pA). F_init_ of RS cells increased steeply compared to those of BS-GCs. The general excitability of both GCs was not different throughout all steps (RS/BS, n = 18/22). **D**: Summary bar graphs for analyses of 1^st^ AP waveform evoked by somatic rheobase current injection into RS and BS cells. Mean values for AP threshold were -34.9 ± 0.5 mV in RS and -36.7 ± 0.7 mV in BS (*p<0.05). For afterhyperpolarization (AHP) amplitudes, 17.1 ± 0.5 mV in RS and 12.0 ± 0.5 mV in BS (***p<0.001). For half-width duration, 0.70 ± 0.01 ms in RS and 0.79 ± 0.01 ms in BS (***p<0.001; RS/BS, n = 18/22). **E**: Input resistance (R_in_; RS, 116.1 ± 9.6 MΩ; BS, 115.3 ± 6.1 MΩ) and resting membrane potential (RMP; RS, -82.1 ± 1.3 mV; BS, -81.2 ± 0.9 mV) were not different between RS and BS. **F:** Proportion of RS and BS cells depends on the GC maturity. BS cells were more frequently found in the group of mature GCs (R_in_ < 200 MΩ) compared to the less mature GC group (R_in_ < 200 MΩ; RS/BS, n = 16/17). **G:** Distributions (*left*) and mean values (*right*) for maximal dendritic branch order in RS (black) and BS (red) cells (RS, 4.5 ± 0.3, n = 8; BS, 5.1 ± 0.1, n = 18, p = 0.10). **H**: Proportion of RS and BS cells along the dorsoventral axis (RS/BS in dorsal, n = 84/148; in ventral, n = 37/8). Error bars indicate S.E.M. *p<0.05. ***P<0.001. n.s., not significant (p>0.05).

To explore whether the bursting behavior is related to morphological properties of GCs, we counted the maximal branch order (MBO) from z-sections of confocal images of biocytin-filled RS-and BS-GCs (Fig. S1). The MBO of majority of mature GCs was five, while cells with MBO higher than five was only found in BS-GCs and that with lower than 4 is only found in RS-GCs (Fig. 1G). The average MBO of BS-GCs was larger than that of RS-GCs, though the difference did not reach statistical significance (RS, 4.5 ± 0.3, n = 8; BS, 5.1 ± 0.1, n = 18, p = 0.1, Mann-Whitney test). We then examined whether the relative proportion of BS- and RS-GCs differs along the hippocampal dorso-ventral axis. We found that BS-GCs were dominant in the dorsal DG, and its proportion was opposite in the ventral DG. Among 232 recorded neurons in dorsal DG, 148 (64%) GCs were identified as BS-GCs, while only 8 (18%) out of 45 GCs were BS-GCs in ventral DG, indicating that the dorsal DG harbors more BS-GCs compared to the ventral DG (Fig. 1H).

### Subthreshold EPSP summation evoked by a single bout of HFS induces NMDA-dependent LTP at LPP-GC synapses

To investigate whether intrinsic firing patterns have any effects on long-term synaptic plasticity, we recorded excitatory postsynaptic potentials (EPSPs) from RS-GCs or BS-GCs by stimulating lateral perforant pathways (LPP) in the presence of PTX (100 μM, a GABA_A_R blocker) and CGP52432 (1 μM, a GABA_B_R blocker) (Fig. 2A). After measuring the baseline EPSPs evoked by stimulation of LPP in a 10 s interval for about 5 min, a single bout of high frequency stimulation (HFS, 10 stimuli at 100 Hz) was applied. For the HFS, we tested two different levels of electrical stimulation intensity: low intensity to induce subthreshold response (HFS_L_) and high intensity to evoke at least 3 APs (HFS_H_). The average stimulation intensities of HFS_L_ and HFS_H_ were 15.6 ± 0.9 V (n = 21) and 25.7 ± 1.45 V (n = 18), respectively (Fig. S2A). The average amplitudes of baseline EPSPs induced by HFS_L_ and HFS_H_ were 5.9 ± 0.3 and 13.8 ± 1.0 mV, respectively (Fig. S2B). Temporal summations of EPSPs evoked by HFS_L_ reached their peaks between -60 mV and -40 mV at the 6^th^ or 7^th^ stimulus. RS- and BS-GCs showed no detectable difference in the temporal summation kinetics (RS, black; BS, red; Fig. 2B). Despite that HFS_L_ of LPP evoked no postsynaptic AP, it induced long-term potentiation (LTP) of EPSP amplitudes in both GC groups, which lasted at least 30 min (Fig. 2C). We denoted this form of LTP as LTP_Sub_, which stands for LTP induced by subthreshold stimulation. The increase in baseline EPSP amplitudes after HFS_L_ was not different between BS-GCs (36.9 ± 8.9%, n = 13) and RS-GCs (35.2 ± 5.3%, n = 12, p = 0.65). The magnitude of LTP_sub_ was correlated with the peak of EPSP summation (r = 0.54, p < 0.001), and significant LTP_Sub_ was induced when the peak was higher than -60 mV (Fig. S2C). To examine the involvement of NMDAR in LTP_Sub_, we tested the effect of APV (50 μM, a NMDAR blocker) on EPSP responses and LTP expression induced by HFS_L_. APV profoundly suppressed the baseline EPSPs as well as EPSP summation (Fig. 2E), and abolished LTP_sub_ (Fig. 2F). These results suggest that NMDA-dependent LTP can be induced at LPP-GC synapses by a single bout of HFS that evokes only a subthreshold voltage response.

**Fig. 2.**
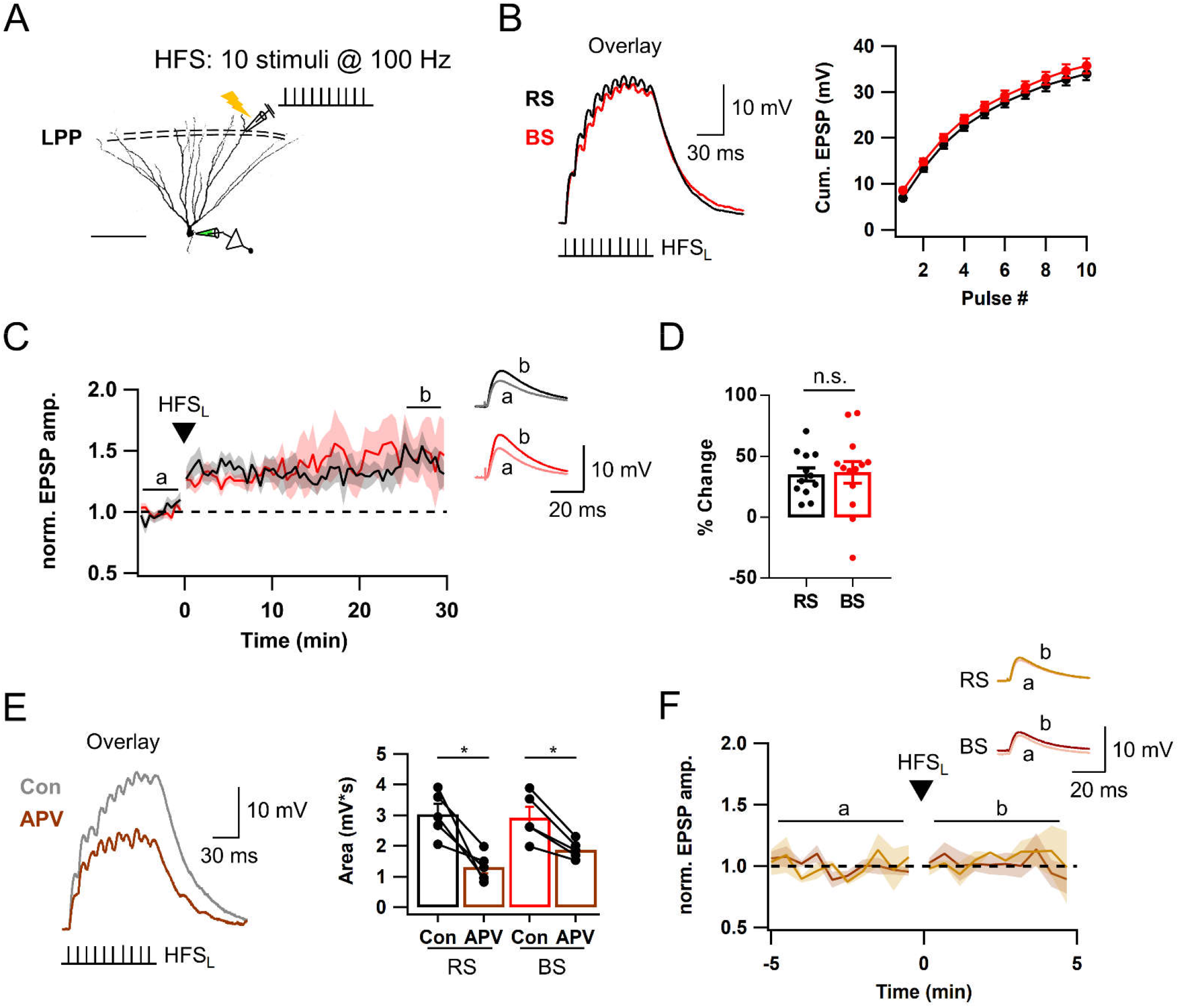
NMDAR-dependent LTP at LPP-GC synapses is induced by a single bout of high frequency stimulation (HFS) at subthreshold level. **A**: Schematic diagram illustrating the recording configuration for synaptic stimulation and whole cell recording of mature GC. Lateral perforant pathway (LPP) in outer molecular layer (OML) was electrically stimulated by a bout of HFS (10 stimuli at 100 Hz). Scale bar is 100 μm. **B**: HFS_L_-evoked subthreshold responses of RS-(black) and BS-GCs (red) (*left*), and their cumulative EPSP amplitudes (*right*). **C**: Time courses of normalized EPSP amplitude before and after HFS_L_ (same as in *B*). Each point represents averaged value for adjacent 3 EPSP amplitudes (30 s binned). Black dashed line denotes baseline EPSP. *Inset*, Representative traces for average of 30 EPSP traces before (a) and 26–30 min (b) after HFS_L_ (This holds for inset traces in all subsequent figures except in *Fig. 2F* and *Fig. 7C*). **D:** LTP magnitudes before and after HFS_L_. There was no significant difference between RS and BS (RS/BS, n = 12/13). **E**: *Left*, Representative traces for EPSP summation in control (black) and after application of APV (brown, 50 μM). *Right*, Mean values for EPSP area in RS [3.0 ± 0.3 mV·s (Con) *vs*. 1.3 ± 0.2 mV·s (APV), n = 5, *p<0.05] and in BS [2.9 ± 0.3 mV·s (Con) *vs*. 1.9 ± 0.1 mV·s (APV), n = 5, *p<0.05]. Note that the APV effect on subthreshold EPSP summation was examined at synapses which have already underwent LTP_sub_. **F**: Time course of normalized EPSP before and after HFS_L_ in the presence of APV in both GCs. EPSP amplitude was not potentiated (RS, 1.7 ± 6.0 %, n = 3, light brown; BS, -2.5 ± 4.5 %, n = 4, brown). *Inset*, EPSPs averaged over 1 to 5 min before (a) and after (b) HFS_L_. Shades and error bars, S.E.M. *p<0.05. n.s., not significant (p>0.05).

### Postsynaptic burst firing is essential for Hebbian LTP at LPP-GC synapses

We then examined whether AP firings in response to HFS_H_ show any difference between BS and RS (Fig. 3A). The F_init_ of HFS_H_-evoked APs was mostly higher than 100 Hz in BS-GCs (128.3 ± 6.9 Hz, n = 21, Fig. 3A). Furthermore, BS-GCs showed a moderate correlation between the F_init_ of synaptically evoked Aps and that of APs evoked by somatic stimulation (r = 0.55, Fig. 3B). In contrast, the F_init_ of HFS_H_-evoked APs in RS-GCs was significantly lower than that in BS-GCs (92.0 ± 9.9 Hz, n = 18; p=0.003; Fig. 3A). These results suggest that mechanisms underlying intrinsic firing pattern contribute to synaptically evoked firing pattern. When the 2nd HFS with higher stimulation intensity (denoted as ‘HFS_H_-2’) was applied 10 min after HFS_L_ by which LTP_Sub_ has been already expressed both in RS and BS, HFS_H_-2 induced further potentiation of EPSPs in BS-GCs, but not in RS-GCs (Fig. 3C). The time course of this LTP induced by HFS_H_-2 is shown as the EPSP amplitudes normalized to the EPSP amplitude just before applying HFS_H_-2 (Fig. 3D). The increase in the EPSP amplitude at 30 min was 44.0 ± 4.8% (n = 7) in BS-GCs, but negligible in RS-GCs (−4.2 ± 7.0%, n=12; p < 0.001). These results indicate that BS-GCs express Hebbian LTP (denoted as LTP_AP_) distinct from NMDA-dependent LTP_Sub_. There was a positive correlation between LTP_AP_ magnitudes and F_init_ of synaptically evoked APs (Fig. 3E). Moreover, when only 1 or 2 APs were elicited by HFS with medium intensity (HFS_M_), LTP was not induced or not maintained even in BS-GCs (−10.6 ± 12.8%, n = 6; Fig. S3A-B), indicating that postsynaptic AP bursts comprised of at least 3 APs at the frequency higher than 100 Hz are essential for the induction of LTP_AP_.

**Fig. 3.**
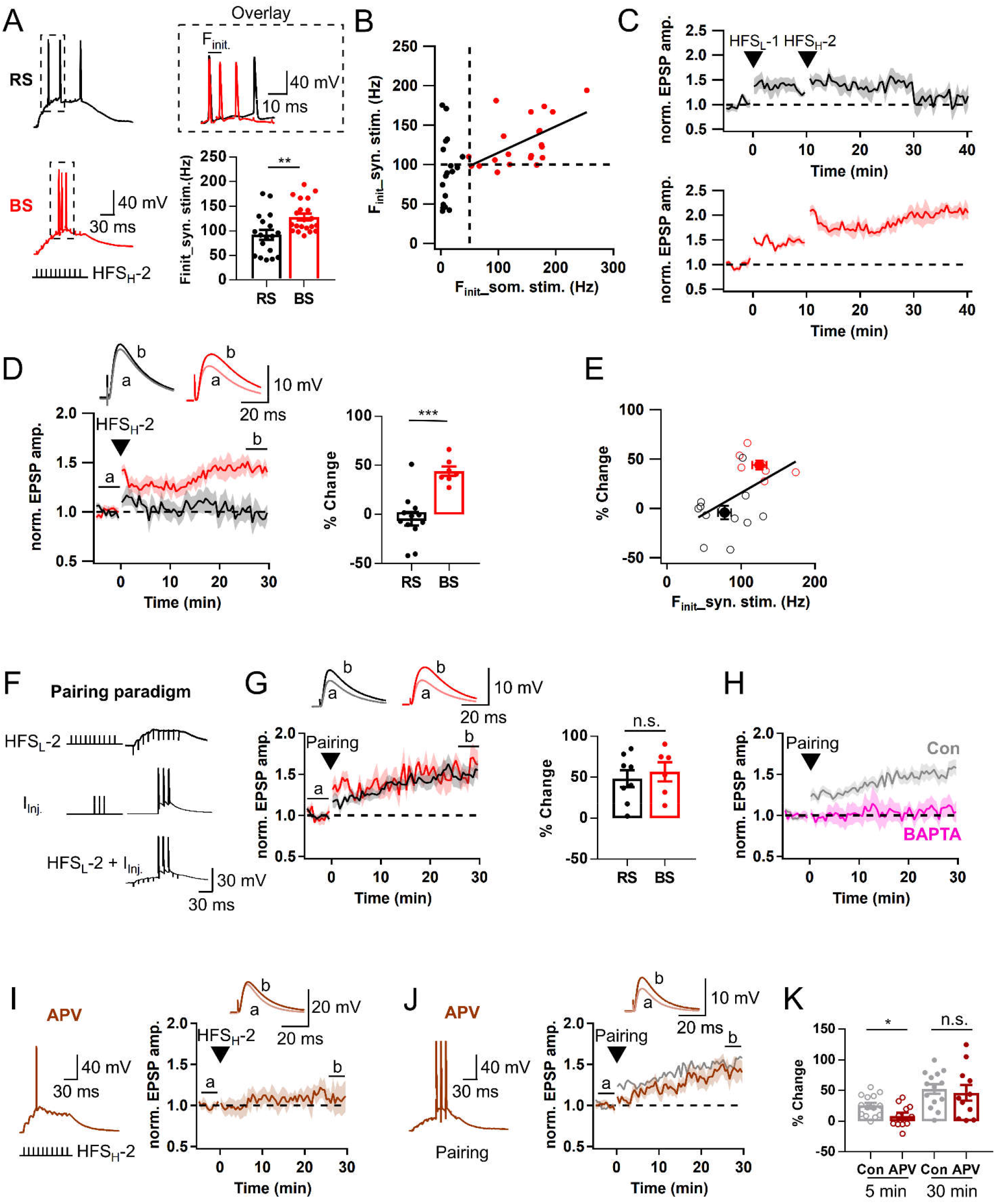
Hebbian LTP depends on post-synaptic AP bursts, and can be induced only in BS. **A**: *Left*, Representative voltage responses in RS (gray) and BS (red) to HFS_H_-2 which elicited 3 APs. *Right upper*, The boxed traces are superimposed for comparison at expanded time scale. *Right lower*, Initial AP frequency (F_init_) of each group (RS/BS, n = 18/21; **p<0.01). **B**. Relationship between F_init_ of APs evoked by somatic current injection and that by synaptic stimulation. Two parameters were significantly correlated in BS (r = 0.55, **p<0.01) but not in RS (r = 0.11). Black bold line, linear regression in BS. r, Pearson’s correlation coefficient. **C**: Time course of normalized EPSP changes induced by applying two sequential HFS (HFS_L_ and HFS_H_) in RS (black, n = 5) and BS (red, n = 5). Note that LTP_AP_ was induced on top of LTP_sub_ in BS, not in RS. **D**: *Left*, Time course before and after HFS_H-_2. *Right*, Magnitude of LTP_AP_ in RS- and BS-GCs (RS: -4.2 ± 7.0%, n = 12; BS, 44.0 ± 4.8%, n = 7, ***p<0.001). **E**: LTP magnitude as a function of synaptically evoked F_init_. F_init_ was correlated to LTP magnitude (r = 0.50, *p<0.05). Open circles, individual data; Closed circles, averaged value for each group. Black line, linear regression line. **F**: Pairing protocol for LTP_AP_ induction. It consists of subthreshold HFS (HFS_L_-2) and post-synaptic 3 APs evoked by somatic pulses (2 ms, 3 nA at 100 Hz). **G**: *Left*, Time course of normalized EPSP before and after a pairing protocol. *Right*, Pairing protocol-induced LTP_AP_ in RS- and BS-GCs (RS/BS, n = 8/6). **H**: LTP was not induced in the presence of intracellular solution containing BAPTA (10 mM, pink, n = 7). The pairing protocol-induced LTP time courses in RS and BS under control conditions shown in *G* were merged, and superimposed in gray (n = 14). **I:** *Left*, Representative voltage traces evoked by HFS_H_-2 in the presence of APV (brown, 50 μM). No AP burst was elicited in the presence of APV. *Right*, Time course of normalized EPSP before and after HFS_H_-2. **J**: Similar as in *I*, but a pairing protocol was applied instead of HFS_H_-2. The control trace (gray) was reproduced from panel H for comparison. **K**: Early (STP, open circle) and late phase (LTP, closed circle) LTP induced by a pairing protocol with and without APV (Control, n = 14; APV, n = 11). Shades and error bars, S.E.M. *p<0.05. **p<0.01. ***p<0.001. n.s., not significant (p>0.05).

To further test the importance of AP frequency for induction of LTP_AP_, we applied a pairing protocol, in which 10 EPSPs were evoked by HFS_L_ coinciding with 3 APs at 100 Hz evoked by brief current injection to the soma (Fig. 3F, see Methods). The pairing protocol successfully induced LTP regardless of cell types with no significant difference in the LTP magnitude between RS-GCs and BS-GCs (Fig. 3G), but 3 APs at 50 Hz failed to induce LTP (Fig. S3C). These findings show that RS-GCs could express LTP_AP_ as if BS-GCs did as long as high frequency APs are paired with synaptic stimulation. Therefore, we did not distinguish BS and RS but pooled the BS and RS data when we analyzed LTP response induced by pairing protocol (gray trace in Fig. 3H). LTP was not induced when intracellular Ca^2+^ was chelated with a high concentration of BAPTA (10 mM, Fig. 3H). These results confirm that at least 3 APs at the frequency higher than 100 Hz are required to activate Hebbian LTP, and suggest that Ca^2+^-dependent mechanisms underlie this form of LTP.

### NMDAR mediates the early phase LTP and facilitates EPSP summation at LPP-GC synapses

To examine whether LTP_AP_ shares the same Ca^2+^ source with NMDAR-dependent LTP_sub_, we tested the effect of APV. Because APV profoundly suppressed EPSP summation (Fig. 2E), in the presence of APV it was difficult to generate 3 APs even with high intensity stimulation, and thus LTP_AP_ was not induced (Fig. 3I), indicating that NMDAR current is critical for EPSP summation to elicit high frequency AP generation. However, we could induce LTP by the pairing protocol in the presence of APV (Fig. 3J). Because the time course of LTP development was distinct from that of control pairing-induced LTP (brown vs. gray traces in Fig. 3J), we compared the LTP magnitudes in the APV conditions with the control values for the early and late phases. To this end, we measured normalized EPSP amplitudes averaged over 1 to 5 min and over 26 to 30 min after HFS, and denoted as short-term potentiation (STP) and and long-term potentiation (LTP), respectively. STP in the APV conditions was significantly lower, while LTP was not different compared to the corresponding control values [STP, 8.6 ± 5.5% vs. 25.7 ± 4.6%, p<0.05; LTP, 46.4 ± 12.7% vs. 52.62 ± 7.71%, p = 0.57; APV (n = 6) vs. Control (n = 14), Mann-Whitney test. Fig. 3K]. These results suggest that the contribution of NMDAR to Hebbian LTP as Ca^2+^ source is limited to the early phase LTP at LPP-GC synapses, whereas it is essential for EPSP summation and AP burst generation.

### T-VDCC contributes to the late phase LTP by facilitating AP bursts at LPP-GC synapses

We showed that burst firing evoked by somatic rheobase current injection (called intrinsic burst firing) has correlation with the F_init_ of synaptically evoked APs which is crucial for LTP_AP_ induction (Fig. 3). We investigated whether ion channel mechanisms underlying intrinsic burst firing also contribute to LTP_AP_. Since T-VDCC is known to mediate intrinsic bursting in DG-GCs (Dumenieu *et al*., 2018), we investigated the role of T-VDCCs in burst firing behavior and LTP_AP_ induction in BS-GCs. Bath application of NiCl_2_ (50 μM, the blocker of T-VDCC) significantly reduced F_init_ of intrinsic burst firing (Control, 171.8 ± 13.3 Hz; NiCl_2_, 38.1 ± 8.89 Hz, n = 9; p<0.01; Wilcoxon signed-rank test; Fig.4A). When the bursts were synaptically evoked, NiCl_2_ partially but significantly reduced the F_init_ (Control, 128.3 ± 6.9 Hz, n = 21; NiCl_2_, 91.2 ± 3.5 Hz, n = 13, p<0.001, Mann-Whitney test; Fig. 4B). Nevertheless, in the presence of 50 μM Ni^2+^, temporal summation of EPSPs evoked by HFS_L_ was little affected (n = 10, p = 0.11; Fig. 4C), and HFS_H_ was able to induce LTP_AP_ in the BS cells (Fig. 4D). In contrast to APV, the early phase LTP was preserved in the presence of Ni^2+^ [STP of Ni^2+^ (n = 7) vs. control (n = 7), 24.9 ± 5.6 vs. 27.5 ± 6.3%, p = 0.902], but no further increase in the EPSP amplitudes was observed (Fig. 4D vs. Fig. 3D), and thus LTP was lower than the control (20.4 ± 7.7 vs. 44.6 ± 5.7%, p<0.05, Fig. 4D). Because Ni^2+^ lowered the F_init_ of synaptically evoked AP bursts, we tested if suppression of late LTP_AP_ can be rescued by pairing protocol. The mean value for LTP measured after the pairing protocol was slightly lower but not significant compared to pairing-induced LTP in control (32.6 ± 13.0 vs. 52.6 ± 7.7%, n = 8, p = 0.19; Fig. 4E), suggesting partial or little contribution of T-VDCC to the LTP_AP_ induction. Similar to HFS_H_-induced LTP, STP was not different from the control value (25.0 ± 8.2 vs. 25.7 ± 4.6%, p = 0.97). These results suggest that T-VDCC primarily contributes to the late phase LTP_AP_ by enhancing F_init_.

**Fig. 4.**
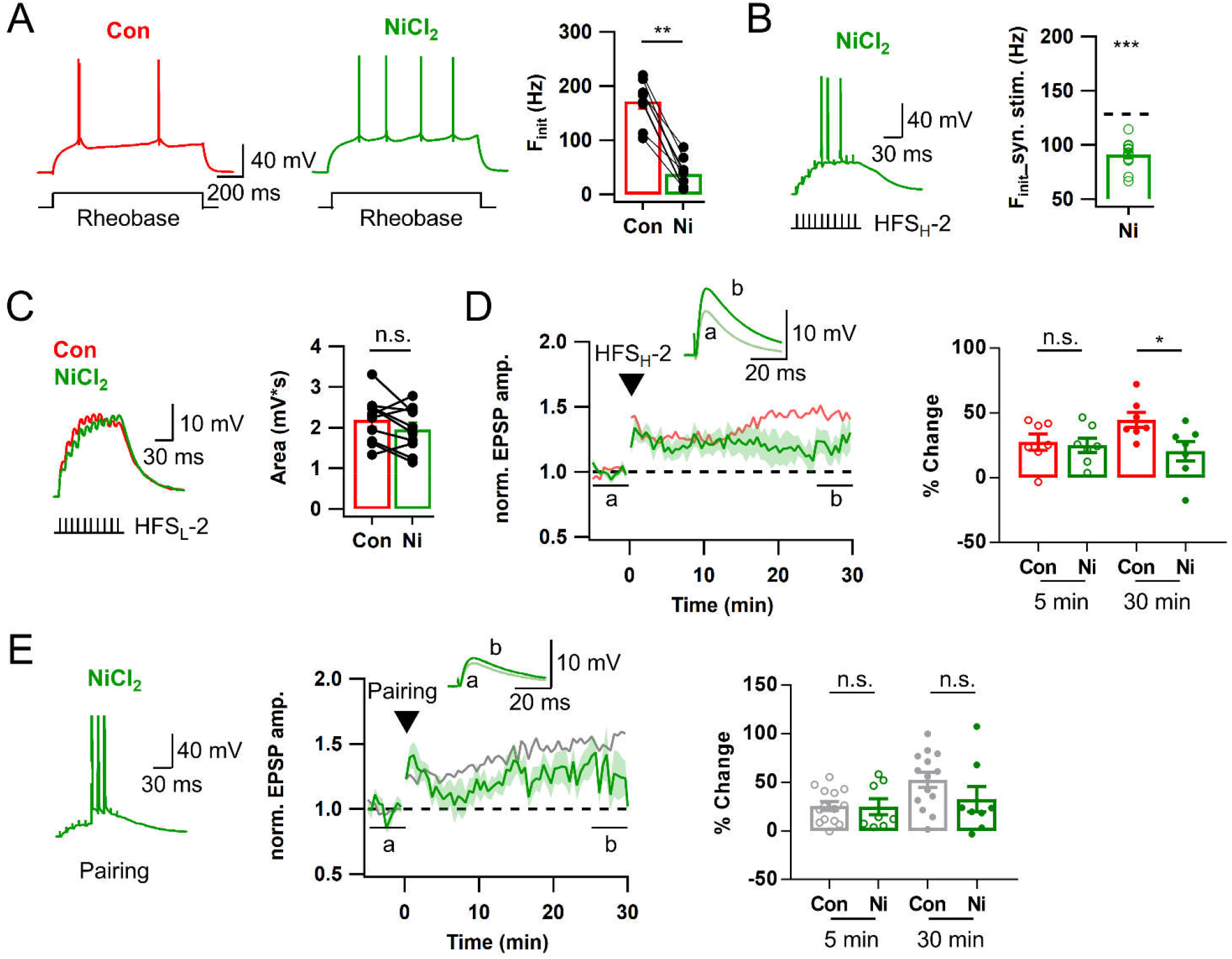
T-type Ca^2+^ channels contributes to the late phase Hebbian LTP by facilitating AP bursts. **A**: *Left & Middle*, Voltage responses of a BS cell to somatic rheobase current injection before (red, Con) and after bath application of NiCl_2_ (50 μM, green). *Right*, Mean values for F_init_ before and after NiCl_2_ application (n = 9). **B**: APs evoked by HFS_H_-2 in a BS cell (*left*) and their F_init_ (*right*) in the presence of NiCl_2_ (n = 13). Black dashed line on the bar graph, control mean F_init_ in BS cells (128.3 Hz). **C:** EPSP summation evoked by HFS_L_-2 (*left*) and the mean area (*right*) before and after applying NiCl_2_ (Con, 2.2 ± 0.2 mV·s; NiCl_2_, 2.0 ± 0.2 mV·s; n = 10). **D**: *Left*, Time courses of normalized EPSP in BS cells before and after HFS_H_-2 with (green) and without (light red) NiCl_2_. The control time course was reproduced from *Fig. 3D* for comparison. *Right*, Magnitude of HFS_H_-induced LTP in the early (STP, open circle) and late (LTP, closed circle) phases in BS cells (Control, n = 7; NiCl_2_, n = 7). **E**: Similar as in *D*, but evoked by a pairing protocol (Control, n = 14, gray; NiCl_2_, n = 8, green). Shades and error bars, S.E.M. *p<0.05. **p<0.01. ***p<0.001. n.s., not significant (p>0.05).

### Persistent Na^+^ current amplifies LPP-evoked EPSP summation and is essential for burst firing

Previously, it was shown that T-VDCC in axon initial segment plays a key role in intrinsic burst firing of GCs (Dumenieu *et al*., 2018). Whereas Ni^2+^ abolished intrinsic bursts (Fig. 4A), it partially reduced F_init_ of synaptically evoked bursts with little effect on EPSP summation (Fig. 4B-C), implying a possible involvement of dendritic channels in synaptically evoked AP bursts. As a candidate ion channel regulating intrinsic and synaptically evoked bursts, we examined persistent sodium current (I_Na.P_). In CA1 pyramidal cells, I_Na.P_ amplifies subthreshold EPSPs leading to spatially tuned firing (Hsu *et al*., 2018). We measured F_init_ of intrinsic bursts in BS-GCs after applying riluzole (10 μM), a typical I_Na.P_ blocker (Chen *et al*., 2005; Yue *et al*., 2005; Hsu *et al*., 2018). Riluzole significantly reduced F_init_ of the intrinsic bursts (Fig. 5A) similar to its effect in CA1 pyramidal neurons (Chen *et al*., 2005). In addition, it markedly suppressed summation of HFS_L_-evoked EPSPs (Fig. 5B). Due to the substantial inhibition of EPSP summation by riluzole, it was not possible to synaptically evoke AP bursts, even with very high stimulation intensity, and LTP was not induced (Fig. 5C). When 10 EPSP bursts induced by HFS_L_ were paired with 3 APs (pairing protocol), however, the late phase LTP was completely rescued (LTP, 43.3 ± 14.7 vs. 52.6 ± 7.7%, n = 10, p = 0.34 compared to pairing-induced LTP in control, Mann-Whitney test; Fig. 5E). The rescue of late phase LTP by the pairing protocol suggests that burst APs coincident with synaptic inputs is essential for the late LTP induction. By contrast, early phase LTP (STP) was significantly lower than the control value (STP, 9.6 ± 6.7 vs. 25.7 ± 4.6%, n = 10, p<0.05), resulting in the LTP time course similar to that in the APV condition (Fig. 3I-K). This similarity may be explained by assuming that Ca^2+^ influx through NMDAR mediate the early phase LTP, and that I_Na.P_ contributes to NMDAR activation in distal dendrites by amplifying EPSP summation, which cannot be compensated by somatic bursts.

**Fig. 5.**
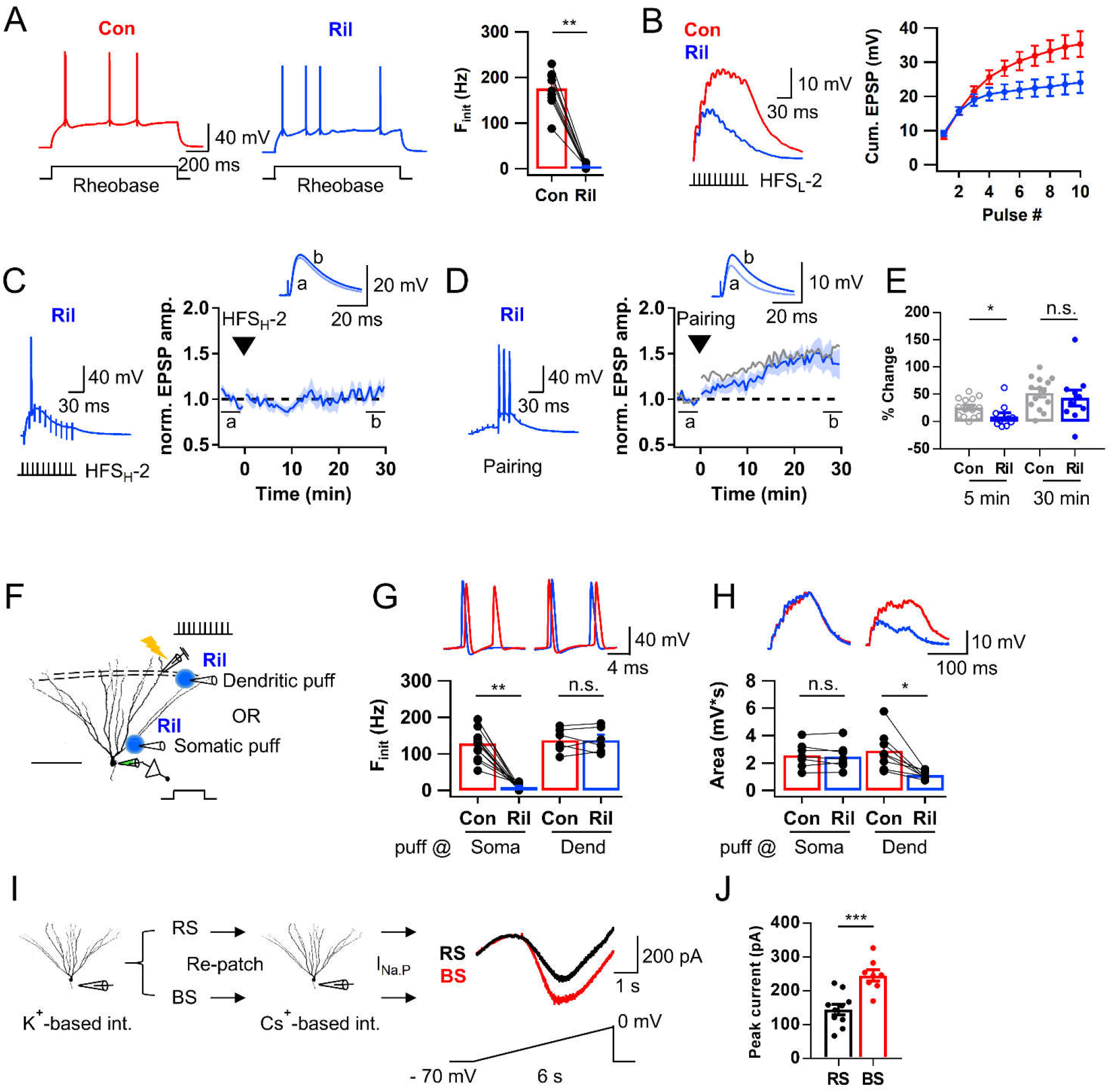
Persistent Na^+^ current (I_Na.P_) amplifies LPP-evoked EPSP summation and is essential for burst firing. **A**: *Left & Middle*, AP trains in BS elicited by somatic rheobase current injection in control (red, Con) and after applying riluzole (blue, Ril, 10 μM). *Right*, Mean F_init_ before and after application of riluzole (Con, 176.5 ± 12.9 Hz; Ril, 5.7 ± 1.4 Hz; n = 10; **p<0.01). **B:** *Left*, EPSP summation evoked by HFS_L_-2 before and after applying riluzole. *Right*, Cumulative EPSP amplitudes in control and riluzole conditions (n = 24). **C:** Representative voltage response to HFS_H_-2 (*left*) and time course of normalized EPSP before and after HFS_H_-2 (*right*, n = 5) in the presence of riluzole. **D**: Similar as in *C*, but evoked by a pairing protocol. **E:** Early (STP, open circle) and late phase (LTP, closed circle) LTP evoked by a pairing protocol in control and riluzole conditions, showing that late LTP_AP_ was rescued (Con, n = 14; Ril, n = 10). **F**: Cartoon for focal application of riluzole (50 μM) at soma or dendrite during somatic current injection or HFS_L_ of LPP. Scale bar is 100 μm. **G:** Representative traces (*upper*) and mean F_init_ (*lower*) of intrinsic AP bursts with somatic (*left*) and dendritic (*right*) puff of riluzole [Soma, 128.8 ± 12.8 Hz (Con) *vs*. 9.0 ± 3.0 Hz (Ril), n = 12, **p<0.01; Dend, 137.5 ± 13.3 Hz (Con) *vs*. 137.6 ± 14.1 Hz, n = 6]. **H:** Similar as in *G*, but area of subthreshold EPSP summation evoked by HFS_L_-2. The EPSP summation was not reduced by somatic puff (*left*) but by dendritic puff (*right*) [Soma, 2.6 ± 0.3 mV·s (Con) *vs*. 2.5 ± 0.4 mV ·s (Ril), n = 7; Dend, 2.9 ± 0.5 mV·s (Con) *vs*. 1.1 ± 0.1 mV·s, n = 8]. **I**: *Left*, Procedure for measuring I_Na.P_ in GCs. *Right*, Representative current responses of RS-(black) and BS-GC (red) to a voltage ramp. **J:** Peak current amplitudes of RS and BS cells (RS, 144.9 ± 15.6 pA, n = 10; BS, 245.4 ± 16.4 pA, n = 8; ***p<0.001). Shades and error bars, S.E.M. *p<0.05. **p<0.01. ***p<0.001. n.s., not significant (p>0.05).

Since riluzole showed profound effects on both intrinsic and synaptically evoked firings, we hypothesized that intrinsic bursting behavior is mainly affected by somatic I_Na,P_, while synaptically evoked AP firings are affected by dendritic I_Na,P_. To test this, we examined the effect of focal puff application of riluzole (Fig. 5F). Peri-somatic puff application of riluzole (50 μM) considerably reduced the F_init_ of intrinsic burst firings in BS-GCs, whereas dendritic puff had no effect at all (Fig. 5G). On the contrary, the EPSP summation was profoundly diminished by dendritic puff, but not by peri-somatic puff of riluzole (Fig. 5H). These results support our hypothesis for the preferential roles of somatic and dendritic I_Na,P_ on the intrinsic and synaptically evoked firing behaviors, respectively. Since BS-GCs have higher F_init_ both for intrinsic and synaptically evoked APs than RS-GCs, we tested whether difference in I_Na.P_ density underlies the different bursting behavior between these two GC types. To measure I_NaP_ in the identified GC type, we first examined the AP responses to somatic rheobase current injection using the standard intracellular solution, carefully withdrew the pipette, and then re-patched the same cell again with Cs^+^-based pipette solution in the presence of Cd^2+^ (200 μM) and TEA (20 mM) in the bath solution to inhibit Ca^2+^ and K^+^ currents (Fig. 5I). We quantified I_Na.P_ in each type of neurons using a slowly rising ramp voltage command protocol from a holding potential of −70 mV to 0 mV for 6 seconds in voltage-clamp configuration. In consistent with our hypothesis, the peak amplitude of I_Na.P_ in BS-GCs was significantly larger than that in RS-GCs (Fig. 5J).

### L-type Ca^2+^ channel is a major Ca^2+^ source for LTP induction but little contributes to firing properties

Above results indicate that NMDAR and T-VDCC partially contribute to LTP_AP_, and that I_Na.P_ is essential for intrinsic and synaptically evoked burst firings in BS-GCs. L-type VDCC (L-VDCC) is known as a calcium source for NMDAR-independent slowly developing LTP induced by 200 Hz tetanic stimuli at CA3-CA1 synapses (Grover & Teyler, 1990; Bayazitov *et al*., 2007). We studied the role of L-VDCC in burst firing and LTP_AP_ induction. Distinct from drugs tested above, nimodipine (10 μM), an L-VDCC blocker, had no effect on the F_init_ of APs evoked by somatic current injection (Fig. 6A). Moreover, nimodipine affected neither EPSP summation induced by HFS_L_ nor the F_init_ of APs evoked by HFS_H_ (Fig. 6B-C). Nevertheless, the late phase of LTP_AP_ was abolished in the presence of nimodipine (LTP, -3.1 ± 11.0 vs. 44.6 ± 5.7%, p<0.01, n = 6, Fig. 6D). Furthermore LTP was not rescued by the pairing protocol (LTP, -2.9 ± 10.3 vs. 52.6 ± 7.7%, p<0.01, n = 7, Fig. 6E), indicating that calcium influx through L-VDCC during AP bursts is essential to induce LTP_AP_. Although the STP values for HFS_H_- and pairing-induced LTP were marginally lowered (HFS_H_, 7.2 ± 9.3%, n = 6, p = 0.10; Pairing protocol, 4.3 ± 8.2%, n = 7, p = 0.08, Mann-Whitney test), the early increase in normalized EPSP was transient, suggesting that it belongs to short-term potentiation (STP) which decayed within 3 min (Lisman, 2017). Therefore, these results indicate that L-VDCC mediate both early and late phase LTP but not STP.

**Fig. 6.**
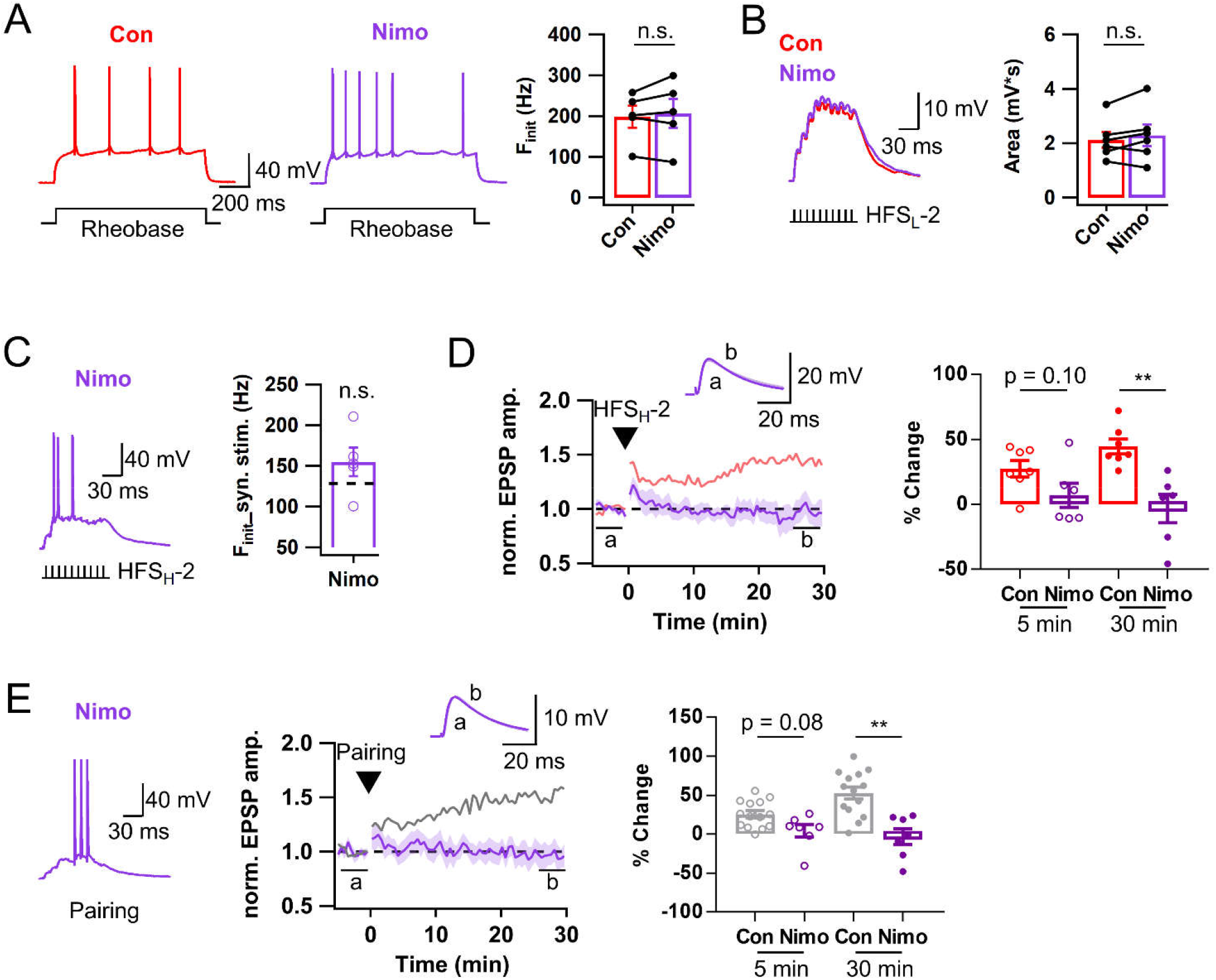
Ca^2+^ influx through L-type Ca^2+^ channels mediates Hebbian LTP at LPP-GC synapses. **A**: *Left & Middle*, Representative AP responses to somatic rheobase current injection before (red, Con) and after application of nimodipine (purple, Nimo, 10 μM). *Right*, F_init_ of intrinsic bursts was not affected by nimodipine (Con, 198.3 ± 26.8 Hz; Nimo, 206.7 ± 36.0 Hz; n = 5). **B:** Exemplar traces (*left*) and mean areas (*right*) of EPSP summation evoked by HFS_L_-2 of LPP before and after applying nimodipine (Con, 2.1 ± 0.3 mV·s; Nimo, 2.3 ± 0.4 mV·s; n = 6). **C**: Exemplar voltage response of BS-GCs (*left*) evoked by HFS_H_-2 and mean F_init_ (*right*) in the presence of nimodipine (Nimo, 154.8 ± 17.5 Hz, n = 5). Black dashed line on the bar graph, control mean F_init_ in BS cells (128.3 Hz). **D**: *Left*, Time course of normalized EPSP in BS-GCs before and after HFS_H_-2. *Right*, Magnitudes of early (STP, open circle) and late phase (LTP, closed circle) LTP in control and nimodipine conditions (Con, n = 7; Nimo, n = 6). The control LTP time course and magnitudes were reproduced from *Fig. 3D* (light red). **E**: Similar as in *C*-*D*, but applied a pairing protocol instead of HFS_H_. The control LTP trace and magnitudes were reproduced from *Fig. 3H* (gray) (Con, n = 14; Nimo, n = 7). Shades and error bars, S.E.M. **p<0.01. n.s., not significant (p>0.05).

### LTP induction at MPP-GC synapses is not affected by firing pattern

Above results indicate that LTP at LPP-GC synapses can be induced by two distinct mechanisms: NMDA-dependent subthreshold LTP and compound Hebbian LTP, and that the latter heavily depends on activation of L-VDCC resulting from postsynaptic AP bursts. We investigated whether MPP-GC synapses share the same LTP mechanisms with those of LPP-GC synapses. We recorded MPP-evoked baseline EPSPs using an electrode placed in the middle of the molecular layer in a 10 s interval for about 5 min before HFS was applied (Fig. 7A). The stimulation intensity was adjusted so that the peaks of EPSP summation evoked by 100 Hz 10 stimuli remained subthreshold level (around -60 mV ∼ -40 mV) (denoted as HFS_L_). Average stimulation intensity of HFS_L_ was 10.9 ± 0.6 V, which is significantly smaller than that used for LTP_sub_ induction at LPP-GC synapses (15.6 ± 0.9 V, Fig. S2A). The EPSP summation usually reached its peak at 2^nd^ or 3^rd^ stimulation and declined afterwards, consistent with the characteristic short-term depression at MPP-GC synapses (Colino & Malenka, 1993). RS- and BS-GCs showed no detectable difference in their subthreshold EPSP responses to HFS_L_ (RS, black; BS, red; Fig. 7B). Unlike NMDAR-dependent LTP_sub_ at LPP-GC synapses, HFS_L_ did not induce LTP either in RS-or BS-GCs [RS, 17.7 ± 8.8% (n = 5); BS, 9.5 ± 7.0% (n = 6); p = 0.66; Fig. 7C]. Moreover, the 1st EPSP amplitude was not affected by APV (MPP, n = 6, p = 0.075; LPP, n = 10, p = 0.047), and the APV effect on the area of EPSP summation was weaker compared to LPP-GC synapses (MPP, p = 0.03; LPP, p < 0.01; Fig. 7D-E). Lower contribution of NMDAR current may be attributable to lower local depolarization at synaptic sites and/or to lower density of NMDAR at the middle part of GC dendrites compared to that at distal dendrites.

**Fig. 7.**
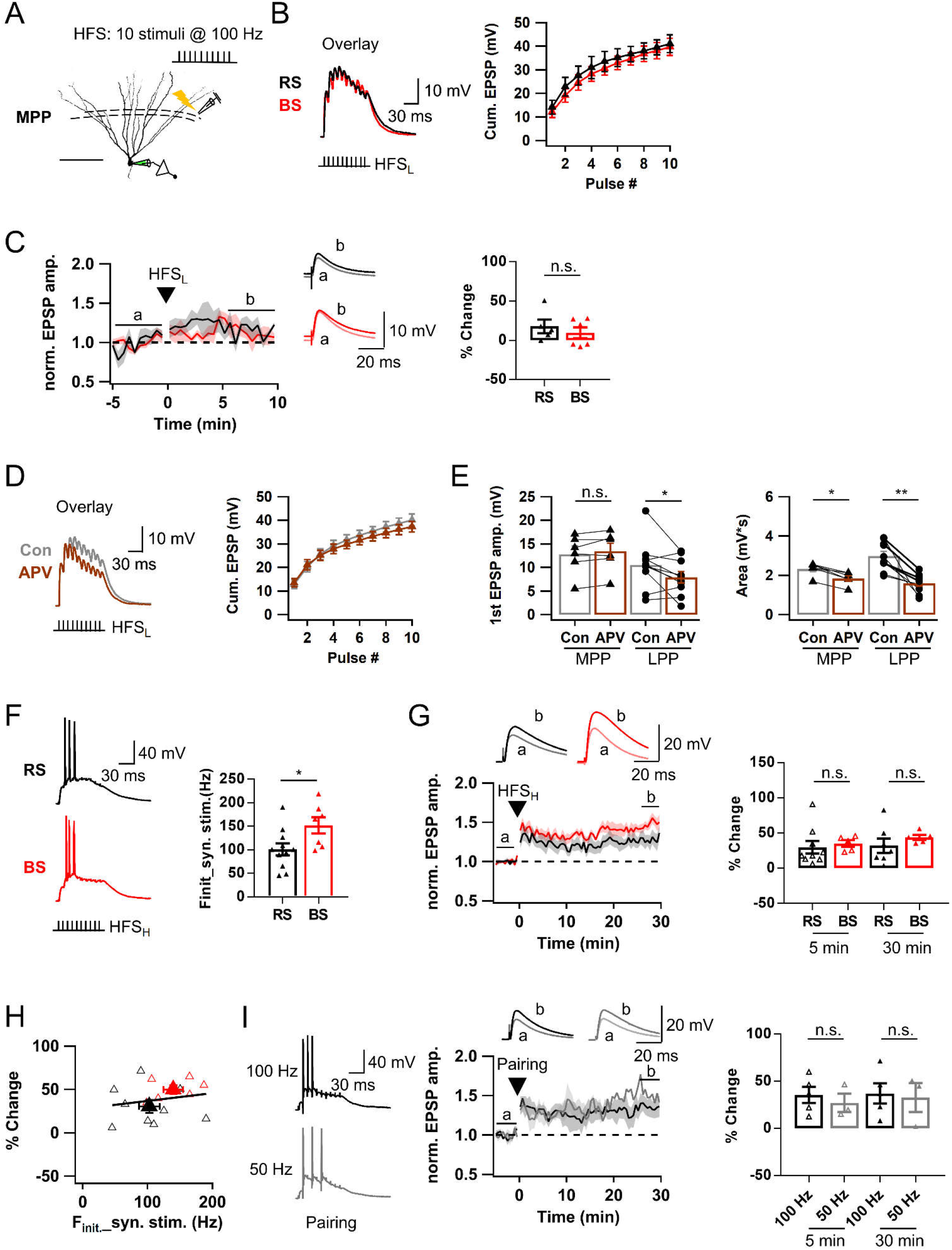
Postsynaptic AP bursts are not required for LTP induction at MPP-GC synapses. **A**: Similar as in *Fig.2A*, but medial perforant pathway (MPP) in medial molecular layer was electrically stimulated with a single bout of train pulses (10 stimuli at 100 Hz). Scale bar is 100 μm. **B**: *Left*, Subthreshold voltage responses evoked by HFS_L_ of MPP in RS-(black) and BS-GCs (red). *Right*, Cumulative EPSP amplitudes of EPSP summation. **C**: *Left*, Time course of normalized EPSP before and after HFS_L_ of MPP. *Right*, Change in normalized EPSP before and after HFS_L_ (RS, 17.7 ± 8.8%, n = 5; BS, 9.5 ± 7.0%, n = 6). **D**: *Left*, Representative EPSP summation in control (black) and after application of APV (50 μM, brown). *Right*, Cumulative EPSP amplitudes in control and APV conditions. **E:** *Left*, Mean amplitude of 1^st^ EPSP evoked by a bout of HFS_L_ in control and APV conditions at MPP and LPP [MPP, 12.8 ± 1.7 mV (Con) *vs*. 13.5 ± 1.7 mV (APV), n = 6; LPP, 9.8 ± 0.9 (Con) *vs*. 7.3 ± 1.1 mV (APV), n = 11, *p < 0.05]. *Right*, Mean area of HFS_L_-induced EPSP summation [MPP, 2.3 ± 0.1 mV·s (Con) *vs*. 1.8 ± 0.1 mV·s (APV), n = 6, *p < 0.05; LPP, 3.0 ± 0.2 mV·s (Con) *vs*. 1.6 ± 0.1 mV·s (APV), n = 10, **p<0.01]. Both plots show weaker contribution of NMDAR to EPSPs at MPP-GCs than LPP-GCs. **F**: *Left*, Representative 3 AP bursts evoked by HFS_H_ in RS-(black) and BS-GCs (red). *Right*, Initial AP frequency (F_init_) of each cell type (RS, 101.2 ± 12.7 Hz, n = 11; BS, 152.1 ± 17.3 Hz, n = 7; *p<0.05). **G**: *Left*, Time course of normalized EPSP before and after HFS_H_. *Right*, Magnitudes of early (STP, open triangle) and late phase (LTP, closed triangle) LTP, showing that no significant difference in HFS_H_-induced LTP_AP_ magnitude between RS- and BS-GCs. **H:** LTP magnitudes as a function of F_init_ of AP bursts evoked by HFS_H_ at MPP-GCs (r = 0.20, p = 0.47). Open symbols, data of individual cells. Closed symbols, averaged value of each group. Black line, linear regression line. r, Pearson’s correlation coefficient. **I:** *Left*, Voltage responses evoked by a pairing protocol comprised of postsynaptic 3 APs at 100 Hz (*upper*, black) or 50 Hz (*lower*, gray) with HFS_L_ of MPP. *Middle*, Time course of normalized EPSP amplitude before and after the pairing protocol. *Right*, Magnitudes of early (STP, open symbols) and late phase (LTP, closed symbols). Note that no difference was found in LTP_AP_ magnitude between pairing at 100 Hz and even at 50 Hz. Shades and error bars, S.E.M. *p<0.05. **p<0.01. n.s., not significant (p>0.05). Shades and error bars, S.E.M. *p<0.05. **p<0.01. n.s., not significant (p>0.05).

When the stimulation intensity was increased to evoke 3 APs, F_init_ was higher in BS-GCs than RS-GCs (Fig. 7F), suggesting that intrinsic bursting mechanisms affect synaptic bursting induced by MPP stimulation, as was shown for LPP-evoked bursts. In spite, LTP_AP_ was induced similarly in both BS and RS [LTP, BS, 43.8 ± 3.6% (n = 5); RS, 32.1 ± 10.2% (n = 7); p = 0.20; Fig. 7F]. Because of no difference between RS and BS in the LTP magnitudes and time courses, the LTP data from the two cell types were merged for following comparison with LTP under different conditions. At MPP-GC synapses, the LTP magnitude was not correlated with F_init_ (Fig. 7G, r = 0.20, p = 0.47), suggesting that AP frequency is not critical for the LTP_AP_ induction at MPP-GC synapses. To further test this idea, we tried to induce LTP using a pairing protocol, in which HFS_L_ was paired with 100 Hz or 50 Hz three APs evoked by somatic stimuli to mimic firing of BS-or RS-GCs, respectively. We found that the LTP magnitudes were not significantly different between them (LTP, 32.6 ± 15.4 vs. 37.1 ± 10.7%, 50 Hz (n = 3) vs. 100 Hz (n = 5), p = 1.00, Fig. 7H).

### Hebbian LTP at MPP-GC synapses is mediated by T-VDCC

We characterized the Ca^2+^ source mediating Hebbian LTP (LTP_AP_) at MPP-GC synapses. Consistent with the small contribution of NMDAR to EPSP summation at MPP synapses (Fig. 7D), AP bursts were readily evoked by HFS_H_ of MPP in the presence of APV (Fig. 8A). In the presence of APV, the late phase LTP magnitude was not different from the control value [LTP, 42.7 ± 11.7 vs. 37.0 ± 6.2%, APV (n = 6) vs. control (n = 12), p = 0.75], but the early phase LTP was significantly inhibited [STP, 8.6 ± 3.1 (n = 7) vs 31.8 ± 5.8% (n = 14), p<0.01] (Fig. 8A), suggesting that NMDAR contributes to short-term potentiation and early phase LTP, but not to the late phase LTP at MPP-GC synapses.

**Fig. 8.**
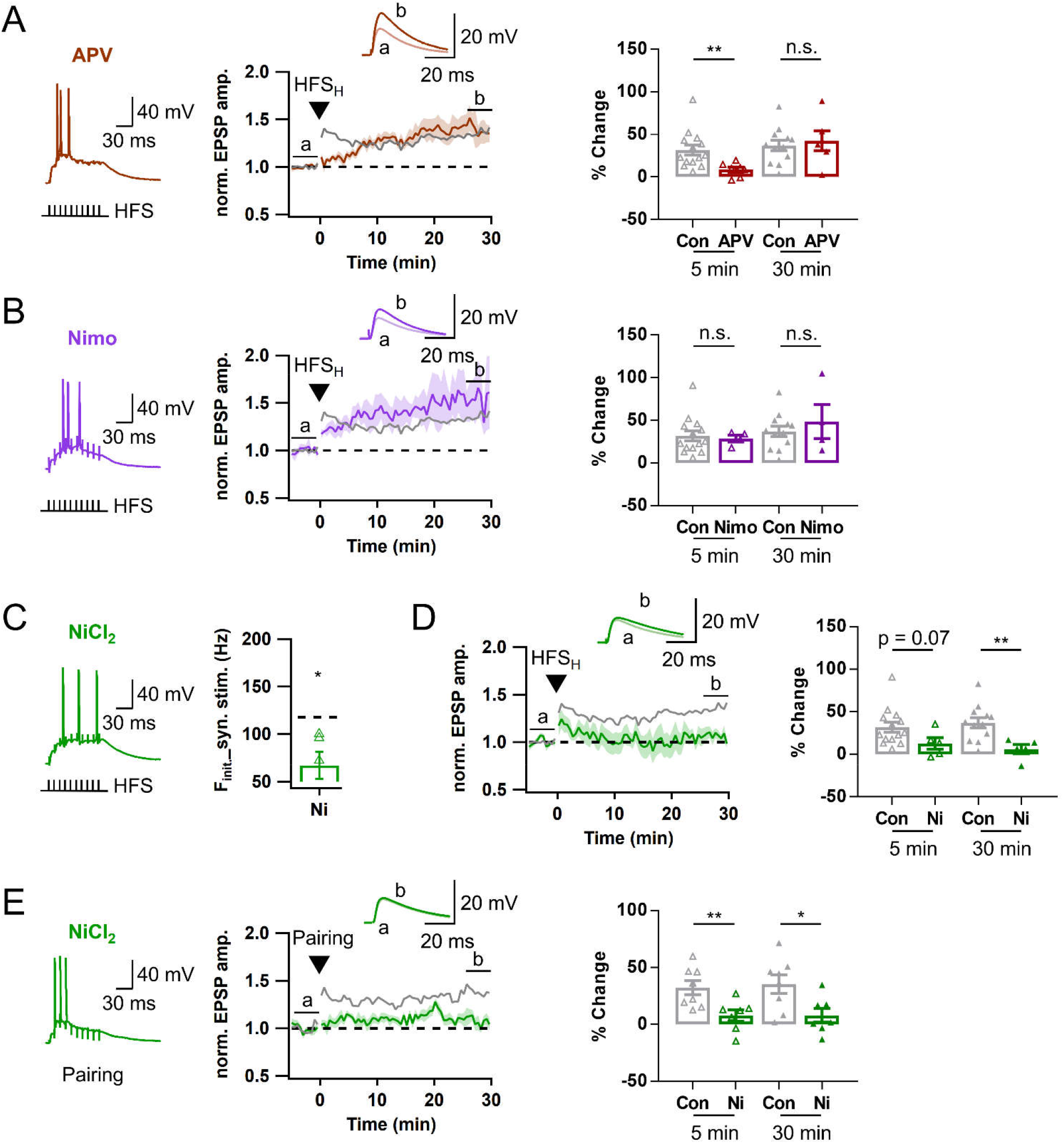
Ca^2+^ influx through T-type is critical to induce Hebbian LTP at MPP-GCs. For comparison, the LTP time courses and LTP magnitudes of RS and BS in *Fig.7G* are merged and shown as a control LTP time course and magnitude (gray). **A**: *Left*, Representative voltage response to HFS_H_ of MPP in the presence of APV (brown, 50 μM). *Middle*, Time course of normalized EPSP before and after HFS_H_. *Right*, Magnitudes of early (STP, open triangle) and late phase (LTP, closed triangle) of LTP, indicating specific reduction of the early phase LTP by APV (Control, n = 12; APV, n = 6). **B:** Similar as in *A*, but in the presence of nimodipine (Nimo, purple, 10 μM, n = 4). Nimodipine had no significant effect on both early and late LTP_AP_. (Control: n = 12; Nimo: n = 4). **C:** *Left*, Representative voltage response to HFS_H_ of MPP in the presence of NiCl_2_ (green, 50 μM). *Right*, Mean F_init_ in the presence of NiCl_2_ (NiCl_2_: 67.0 ± 14.4 Hz, n = 5). Black dashed line denotes the mean value for F_init_ of both GC types (117.6 Hz, n = 15). **D:** Similar as in *A*, but in the presence of NiCl_2_ (green). Late phase of LTP was significantly inhibited. **E:** Similar as in *D*, but a pairing protocol was applied of HFS_H_. Both early and late LTP is significantly reduced. Shades and error bars, S.E.M. *p<0.05. **p<0.01. n.s., not significant (p>0.05).

Nimodipine had no significant effect on MPP-evoked AP generation similar to LPP synapses. In stark contrast to LTP_AP_ at LPP-GC synapses (Fig. 6), LTP_AP_ at MPP-GC synapses was not affected by nimodipine (STP, 28.7 ± 4.0%, p = 0.95; LTP, 48.5 ± 20.0%, n = 4, p = 0.77; Fig. 8B), but abolished by NiCl_2_. NiCl_2_ significantly reduced the F_init_ of MPP-evoked APs (Fig. 8C), and abolished the late phase LTP at MPP-GC synapses (LTP, 6.0 ± 5.3%, n =5, p = 0.009, Fig. 8D). NiCl_2_ marginally lowered STP (12.8 ± 6.8%, n = 5, p = 0.07), but the early increase in normalized EPSP was not sustained (Fig. 8D), reminiscent of the nimodipine effects at LPP synapses (Fig. 6D). Moreover, the pairing protocol did not rescue the Ni^2+^ effect on LTP_AP_ at MPP-GC synapses [Pairing protocol-induced LTP; STP, 7.8 ± 5.1 vs. 30.3 ± 6.7%, p = 0.017; LTP, 8.0 ± 6.0 vs. 35.4 ± 8.2%, Ni^2+^ (n = 7) vs. control (n = 7), Fig. 8E], indicating that LTP_AP_ at MPP-GC synapse is mediated by Ca^2+^ influx through T-VDCC.

### HFS_H_ activates mGluR5 signaling pathways at LPP-GC synapses

Pairing presynaptic 10 stimuli at lower frequency (50 Hz) with 3 postsynaptic APs at LPP-GC synapses failed to bring significant potentiation (Fig. S3D). Moreover, LTP_AP_ was not induced by 1 Hz repeated pairing of a single presynaptic stimulation with postsynaptic AP bursts for 5 min (pre- and post-synaptic sequence, 5 ms interval; Fig. S4), indicating that LTP_AP_ at LPP-GC synapses critically depended not only on postsynaptic but also on presynaptic bursts. The requirement of presynaptic bursts is consistent with the condition for spillover of synaptically released glutamate thereby peri-synaptic mGluRs could be activated (Okubo & Iino, 2011). For studying downstream signaling of LTP_AP_, we adopted the pairing protocol to avoid possibility that the test drugs may affect postsynaptic AP bursts. In the presence of MPEP (25 μM), the paring protocol did not induce LTP at LPP-GC synapses (Fig. 9A).

**Fig. 9.**
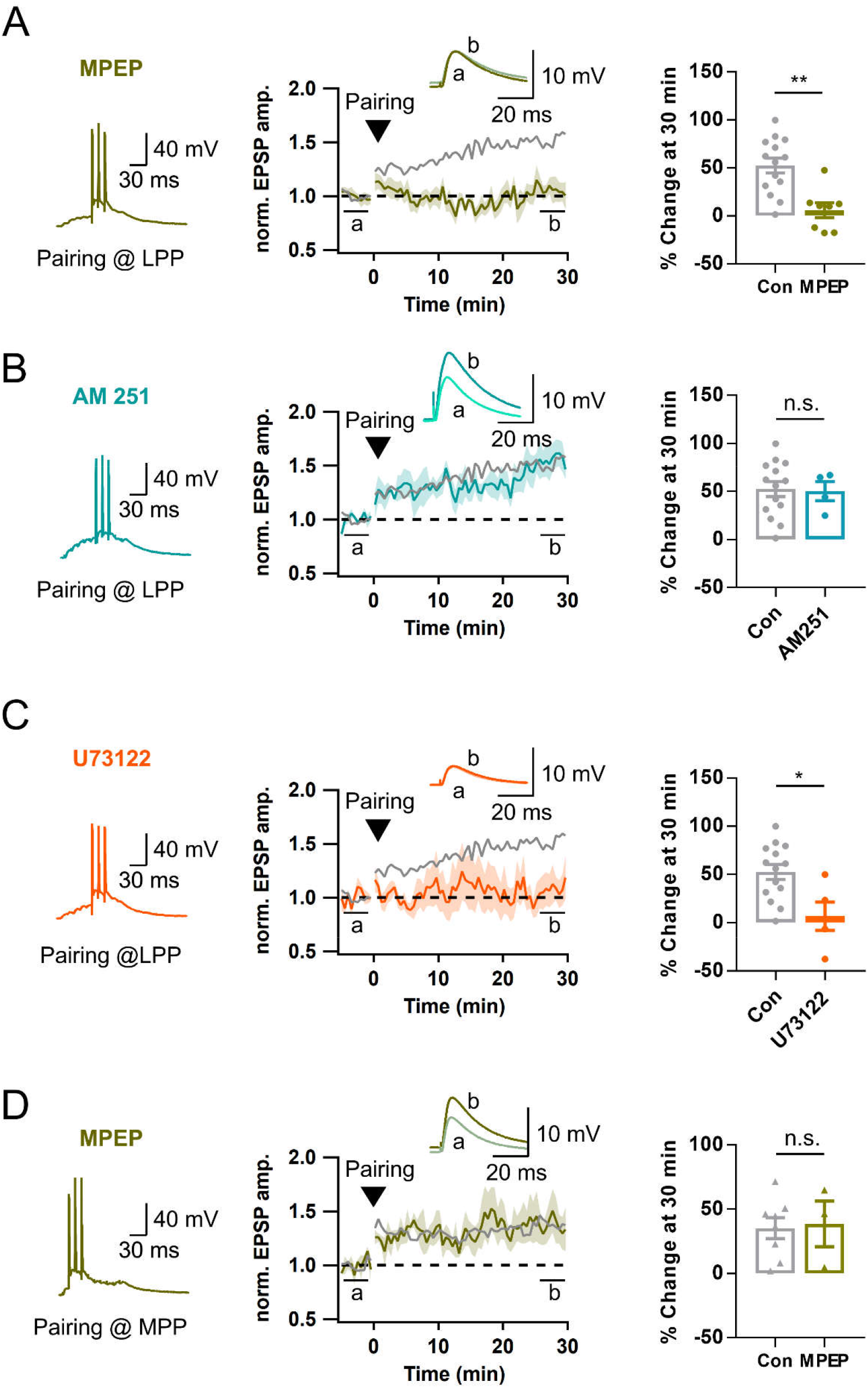
HFS_H_ activates mGluR5 signaling pathways at LPP-GCs, but not at MPP-GCs. **A**: *Left*, Representative voltage response to a pairing protocol at LPP-GC synapse in the presence of MPEP (dark green, 25 μM). *Middle*, Time course of normalized EPSP before and after applying a pairing protocol. *Right*, Magnitudes late phase (LTP, closed circle) of LTP in the presence of MPEP [6.0 ± 7.7% *vs*. 52.6 ± 7.7%, MPEP (n = 8) *vs*. Con (n = 14), **p<0.01]. **B:** Similar as in *A*, but in the presence of AM 251 (light blue, 5 μM). Late LTP_AP_ were not affected by AM251 [50.3 ± 9.9% *vs*. 52.6 ± 7.7%; AM 251 (n = 4) *vs*. Con (n = 14)]. **C:** Similar as in *A*, but in the presence of U73122 (orange, 2 μM). Late LTP_AP_ were significantly inhibited [6.7 ± 14.6% *vs*. 52.6 ± 7.7%, U73122 (n = 5) *vs*. Con (n = 14), *p<0.05]. For comparison of LTP at LPP synapses in *A-C*, the time courses and magnitudes of LTP induced by a pairing protocol are reproduced from *Fig. 3H* (gray). **D:** Similar as in *A*, but at MPP-GCs. The data for LTP induced by 50 and 100 Hz pairing protocol shown in *Fig. 7I* were merged and reproduced as control LTP [38.7 ± 18.8% *vs*. 35.4 ± 8.2%; MPEP (n = 3) *vs*. Con (n = 8)]. Shades and error bars, S.E.M. *p<0.05. **p<0.01. n.s., not significant (p>0.05).

Recently, it was shown that two trains of HFS (1 s at 100 Hz, 1 min interval) of LPP induced presynaptic LTP through activation of mGluR5 and endocannabinoid-dependent retrograde signaling (Wang *et al*., 2016). We tested if LTP_AP_ observed in the present study shares the same mechanism with the LTP form reported in Wang et al (2016). We could induce LTP at LPP-GC synapses by the pairing protocol even in the presence of AM251, a CB_1_ inverse agonist (Fig. 9B), arguing against involvement of endocannabinoid signaling in the induction of LTP_AP_ at LPP-GC synapses.

Because mGluR5 is a G_q_-coupled G protein receptor, we tested involvement of phospholipase C (PLC) in the downstream signaling for induction of LTP. After pre-incubation of the slice with U-73122 (an inhibitor of PLC, 2 μM) at least for 30 min, LTP_AP_ was abolished (Fig. 9B). In contrast to LTP_AP_ at LPP-GC synapses, that at MPP-GCs was not affected by MPEP (Fig. 9C). This result is consistent with a previous report that more intense stimulation is required for induction of mGluR-dependent LTP at these synapses (Wu *et al*., 2008).

## Discussion

### Ionic mechanisms underlying Hebbian LTP at LPP and MPP synapses

One of main findings of the present study is that induction of Hebbian LTP (LTP_AP_) at LPP-GC synapses is critically dependent on high frequency burst firing of pre- and post-synaptic cells. Induction of Hebbian LTP required at least three post-synaptic APs firing at 100 Hz or higher frequency, and thus Hebbian LTP at LPP-GC synapses occurred preferentially at BS-GCs compared to RS-GCs. To scrutinize the mechanisms underlying LTP_AP_ at LPP-BS synapses, we differentiated whether different inward currents contribute to LTP_AP_ by enhancing postsynaptic AP bursts (burst-enhancer) and/or providing Ca^2+^ influx mediating LTP_AP_ (LTP-mediator). To this end, when an inward current blocker suppressed HFS_H_-induced LTP_AP_, we tried to induce LTP_AP_ by applying the pairing protocol in the presence of the blocker. When LTP_AP_ was rescued by the pairing protocol, we regarded it as ‘burst-enhancer’, and otherwise as ‘LTP-mediator’. The other factor to be considered was the time course of LTP_AP_. As shown in Fig. 3D and G, LTP_AP_ in BS-GCs was comprised of three components: immediate, early and late potentiation. The immediate phase decayed within 3 min, and early and late phases of sustained potentiation lasted more than 30 min (Lisman, 2017). Because the mechanism underlying immediate and early phase potentiation are known to be different from that underlying late phase LTP at CA3-CA1 synapses (Grover & Teyler, 1990; Bayazitov *et al*., 2007), we measured normalized EPSP amplitudes averaged over early (1 - 5min) and late (26 - 30 min) intervals of the LTP time course, and regarded the former (STP) and the latter (LTP) as magnitudes of immediate plus early phase potentiation and late phase potentiation, respectively. We examined contributions of NMDAR, T-VDCC, I_Na,p_ and L-VDCC under this framework. For the late phase LTP_AP_ at LPP-GC synapses, only L-VDCC met the condition for the ‘LTP-mediator’, and other inward currents seem to contribute as a burst-enhancer. For the early phase LTP, the LTP magnitude induced by the pairing protocol was marginally or significantly lower in the presence of blockers of NMDA-R and L-VDCC, implying that these two Ca^2+^-influx channels may mediate the early phase LTP. The differential involvements of NMDA-R and L-VDCC in early and late LTP has been shown in CA3-CA1 synapses (Grover & Teyler, 1990). Whereas NMDA-dependent LTP was rapidly expressed in the postsynaptic locus, NMDAR-independent LTP developed more slowly, depended on L-VDCC and expressed in presynaptic locus (Bayazitov *et al*., 2007). These ionic mechanisms of LTP at CA3-CA1 synapses are different from those at LPP-GC synapses, in that L-VDCC contributes to both early and late LTP, while NMDAR does to early LTP. The locus of LTP expression at LPP-GC synapses remains to be investigated. In contrast to LPP-GC synapses, Hebbian LTP at MPP-GC synapses was mediated by T-VDCCs, and BS-GCs had no privilege for induction of LTP at MPP synapses. The requirement of T-VDCC is consistent with (Dumenieu *et al*., 2018), which showed that deletion of CaV3.2 gene reduced LTP at MPP-GC synapses.

### Ionic mechanisms underlying AP bursts

Most neuronal burst firings are associated with prominent afterdepolarization (ADP), which can be generated by dendritic Ca^2+^ spikes and/or axo-somatic slow activating inward current. The dendritic contributions to burst firing has been found in hippocampal and neocortical pyramidal neurons (Larkum *et al*., 1999; Chen *et al*., 2005; Raus Balind *et al*., 2019). The burst firing of GCs seems to be axo-somatic type, because axo-somatic T-VDCCs played a crucial role (Dumenieu *et al*., 2018). We found that not only T-VDCCs but also I_Na,P_ contribute to the burst firings in GCs (Fig. 4 and 5). I_Na,p_ is a small fraction of Na^+^ current that slowly inactivates and exhibits low threshold for activation compared to larger fast and transient fraction of Na^+^ current. It has been suggested that I_Na,p_ not only generates ADP (Yue *et al*., 2005) but also amplifies synaptic current (Schwindt & Crill, 1995). It is very likely that contribution of I_Na,p_ to ADP underlies intrinsic burst firing, while I_Na,p_ contributes to synaptically evoked AP by amplifying EPSP summation. Consistent with this view, we showed that the F_init_ of intrinsic bursts in BS-GCs was reduced by local puff of riluzole to the soma, but not by that to the dendrites, suggesting contribution of somatic I_Na,p_. By contrast, dendritic I_Na,P_, but not somatic I_Na,P_, was responsible for enhancing EPSP summation and LPP-evoked AP bursts (Fig. 5G-H). On the other hand, block of T-VDCC using NiCl_2_ resulted in only partial reduction of F_init_ of LPP-evoked AP bursts (Fig. 4C), whereas it had stronger effect on MPP-evoked burst firing (Fig. 8C), implying higher density expression of T-VDCC on proximal dendrites compared to distal dendrites. This view is supported by our findings that T-VDCC plays as an LTP-mediator in MPP synapses whereas it plays only a partial role in LTP_AP_ induction at LPP synapses (Fig. 4D-E and Fig. 8D-E).

### The role of post-synaptic high frequency bursts in Hebbian LTP at LPP synapses

At LPP-GC synapses, LTP_AP_ was induced preferentially in BS cells and the F_init_ of synaptically evoked burst firing was highly correlated with the LTP_AP_ magnitude (Fig. 3E). Why high frequency bursts are required for LTP_AP_ induction at LPP-GC synapses? L-VDCC was a major Ca^2+^ source mediating LTP_AP_ (Fig. 6D), although L-VDCC had little influence on both synaptic and AP responses (Fig. 6A-B). In light of these results, it is likely that activation of L-VDCC requires high frequency back-propagating APs (bAPs). A previous study in L5 neocortical pyramidal neurons may provide a hint for addressing the question. (Larkum *et al*., 1999), using dual patch recordings at apical dendrite and soma in a single neocortical L5 pyramidal neuron, discovered nonlinear summation of bAPs at distal apical dendrites: As somatic APs back-propagated along apical dendrites, they were attenuated in amplitude and broadened in width. While low frequency bAPs underwent only such linear attenuation, as the bAP frequency increased above a critical point (100 Hz), bursts of four bAPs summated to readily reach the threshold for activation dendritic Ca^2+^ channels (Larkum *et al*., 1999). We imagine that a similar scenario may be involved in the L-VDCC dependent dendritic Ca^2+^ signaling evoked by a burst of three somatic APs in GCs. The broadening and attenuation of bAPs at intermediate dendrites has been shown in our previous study (Kim *et al*., 2018). Summation of bAPs at distal dendrites remains to be elucidated in GCs, though it would be a challenging task considering the feasibility of patching on the distal dendrites of GCs.

### Hebbian vs. non-Hebbian LTP at LPP-GC synapses

Previously we showed a different form of LTP at LPP-GC synapse, which was critically dependent on dendritic Na^+^ spikes and activation of NMDA receptors (Kim *et al*., 2018). This form of LTP was induced by theta burst synaptic stimulation (TBS) of LPP, but not by the standard spike time-dependent plasticity (STDP) protocol, which is pairing EPSP with a single somatic AP (Kim *et al*., 2018). The LTP shown in (Kim *et al*., 2018) could be induced even without somatic APs as long as TBS elicited dendritic spikes. In contrast, a burst of 100 Hz three APs was required for the induction of LTP_AP_ not only in the pairing protocol but also in the induction protocol of synaptic stimulation alone (HFS_H_). Therefore, the LTP forms shown in our previous and present studies belong to non-Hebbian and Hebbian LTP, respectively. The strong attenuation of back-propagating somatic APs along the dendrites of mature GCs might be responsible for the no LTP induction by the standard STDP protocol (Kim *et al*., 2018). The results of the present study suggest that postsynaptic AP bursts may overcome the strong dendritic attenuation probably by summation of bAPs in distal dendrites to activate L-VDCCs.

Remarkably, only a single bout of HFS (10 stimuli at 100 Hz) was sufficient for induction of Hebbian LTP. We previously used TBS (4 repeats of 5Hz 10 bouts of HFS) for induction of dendritic spike-dependent LTP in (Kim *et al*., 2018). The key differences in the LTP induction protocols between these two studies are not only the number of HFS bouts but also the LPP stimulation intensity. The baseline EPSP in our previous and present studies were 7.1 ± 0.5 mV and 13.8 ± 1.0 mV, respectively, indicating that the stimulation intensity required for LTP_AP_ is stronger than that for dendritic spike-dependent LTP.

The subthreshold LTP discovered in the present study has not been described before. It is unique in that ten stimuli which evoked only subthreshold EPSP summation can induce NMDAR-dependent LTP as long as the peak EPSP summation was higher than -60 mV. Because such weak stimuli have been routinely employed to characterize the baseline properties of synapses, the subthreshold LTP has been ignored in our previous study (Kim *et al*., 2018). Therefore, the dendritic spike-dependent LTP described in (Kim *et al*., 2018) has been induced on the top of subthreshold LTP. Given that subthreshold LTP is mediated by NMDAR, the local EPSP summation elicited by high frequency LPP inputs may result in large local depolarization at distal dendrites sufficient for activation of NMDARs, even if it does not elicit somatic APs or dendritic spikes (Note that there was no evidence for dendritic spikes in somatic recordings during the subthreshold LTP induction). Recent in vivo whole-cell recordings in GCs revealed that majority of GCs were under the influence of spatially tuned PP synaptic inputs while only minority of them exhibited spatially tuned firings (Zhang *et al*., 2020). It is not clear whether nonspatial LPP inputs are also extensive similar to such spatial inputs. If it is so, subthreshold LTP at LPP-GC synapses might extensively occur over the GC population receiving brief bursts of LPP inputs independent of postsynaptic firings.

## Materials and Methods

### Slice preparation and electrophysiology

Acute hippocampal slices (thickness, 350 μm) were prepared from the brains of 17-to 25-day-old Sprague-Dawley rats of either sex. Rats were anesthetized (isoflurane, Forane; Abbott) and decapitated immediately. All the experiments were approved by the University Committee Animal Resource in Seoul National University (Approval #: SNU-210825–6). All brains were obtained coronally for dorsal hippocampus or horizontally for ventral hippocampus (coronal sections located between 4.2 mm and 5.6 mm from the posterior end and transverse sections located 2.8 mm and 4.2 mm from ventral end of the right hemisphere). Slices were prepared in an oxygenated ice-cold sucrose-containing physiological saline using a vibratome (VT1200, Leica), incubated at ∼36°C for 30 min, and subsequently maintained in the same solution at room temperature until the recordings. Recordings were performed at near-physiological temperature (33–35°C) in an oxygenated artificial cerebral spinal fluid (ACSF).

Patch pipettes were obtained from borosilicate glass capillaries (outer diameter = 1.5 mm, inner diameter = 1.05 mm) with a horizontal pipette puller (P-97, Sutter Instruments). The open-tip resistance of patch pipettes was 2.5–4.5 MΩ for somatic recordings. Current-or voltage clamp recordings were performed with an EPC-10 USB Double amplifier (HEKA Elektronik). In current-clamp recordings, series resistance was 8–20 MΩ. Pulse protocols were generated, and signals were low-pass filtered at 3 or 10 kHz (Bessel), digitized (sampling rate: 20 kHz) and stored using Patchmaster software running on a PC under Window 10. Resting membrane potential (RMP) was measured immediately after patch break-in. Input Resistance (R_in_) was determined by applying Ohm’s law to the steady-state voltage difference resulting from a hyperpolarizing current step (−20 pA, 500 ms). Threshold for action potential was determined at points at which the derivative of voltage exceeded 40 V/s of somatic or synaptic stimulations. Pipette capacitance and series resistance compensation (bridge balance) were used throughout current-clamp recordings. Recordings were stopped and discarded if the RMP depolarized above -70 mV or hyperpolarized under -90 mV, or if the resting membrane potential or R_in_ changed by more than 20% during the data acquisition.

All experiments were performed on visually identified mature GCs on the basis of the relatively large and round-shaped somata under DIC optics. GCs located at the superficial side of the GC layer in the suprapyramidal blade were purposely targeted. These cells had the average RMP of -81.6 ± 0.7 mV and R_in_ of 115.7 ± 5.5 MΩ, that are similar to characteristic intrinsic properties of mature GC population (Schmidt-Hieber *et al*., 2004). Cells were filled with a fluorescent dye, Alexa Fluor 488 (50 μM, Invitrogen) at least 5 min and imaged with LED system (Thorlabs) mounted on an upright microscope equipped with a 60x water immersion objective lens (N.A. 1.0). In order for focal electrical stimulation (100 μs pulses of 5–40 V intensities) of the medial or lateral perforant pathways, a ACSF-filled glass pipette microelectrode (3–4 MΩ) was placed in the vicinity of intermediate or distal part of a visually identified dendrite (typically at <50 μm distance) of a GC under whole-cell patch. For evaluation of baseline synaptic responses, excitatory postsynaptic potentials (EPSPs) were evoked by applying a pulse every 10 s through a stimulation electrode. All experiments were performed in the presence of the GABA receptor antagonist picrotoxin (PTX, 100 μM) and CGP52432 (1 μM).

### Stimulation protocols for the induction of long-term potentiation (LTP)

LTP was induced by either single bout of high-frequency stimulation (Remy & Spruston, 2007) of afferent fibers or a paring protocol. HFS consists of 10 stimuli at 100 Hz under current clamp mode. Depending on the stimulation intensity, HFS evoked subthreshold EPSP summation alone or additively post-synaptic APs, which are denoted as HFS_L_ and HFS_H_, respectively. The pairing protocol is comprised of HFS followed by post-synaptic injection of three suprathreshold current pulses (2 ms, 3 nA) at 100 Hz with a time delay (50 ms), similar to a protocol in (Watanabe *et al*., 2002). The time delay, 50 ms, was set based on the averaged synaptically evoked firing onset time. For LTP experiments, we monitored baseline EPSPs every 10 s at least for 5 min before applying LTP induction, after which we resumed the EPSP monitoring at least for 30 min. For off-line analysis, EPSP amplitudes were normalized to the mean of baseline values. A time course of normalized EPSP amplitudes was subject to binomial smoothing using a built-in function of IgorPro7 (WaveMetrics). The magnitude of EPSP potentiation was evaluated as a mean of smoothed EPSP amplitudes measured 1 to 5 min or 26 to 30 min after LTP induction [denoted as short-term potentiation (STP) and long-term potentiation (LTP), respectively).

### Solutions and chemicals

The extracellular solution for dissection and storage of brain slices was sucrose-based solution (87 mM NaCl, 25 mM NaHCO_3_, 2.5 mM KCl, 1.25 mM NaH_2_PO_4_,7 mM MgCl_2_, 0.5 mM CaCl2, 10 mM glucose, and 75 mM sucrose). Physiological saline for experiments was standard ACSF (125 mM NaCl, 25 mM, NaHCO_3_, 2.5 mM KCl, 1.25 mM NaH_2_PO_4_, 1 mM MgCl_2_, 2 mM CaCl_2_, and 25 mM glucose).

For whole-cell recording, we used K^+^ rich intracellular solution that contained 115 mM K-gluconate, 20 mM KCl, 10 mM HEPES, 0.1 mM EGTA, 4 mM MgATP, 10 mM Na2-phosphocreatine, and 0.3 mM NaGTP, pH adjusted to 7.2–3 with KOH (∼300 mOsm). If necessary, 50 μM Alexa 488 were added to the internal solution to detect the dendrites. In subset of experiments for measuring persistent sodium current (I_Na.P_), aCSF containing 20 mM tetraethylammonium chloride (TEA) and 0.2 mM CdCl_2_ was used, and an internal solution in which K-gluconate and KCl were replaced with Cs-methanesulfonate and CsCl, respectively, at the same concentration.

### Immunohistochemistry and morphological analysis

GCs were filled with 0.2% biocytin (wt/vol) at least 20 min during whole-cell recording. The acute slices (thickness, 350 μm) were fixed overnight at 4°C in 4% paraformaldehyde (Fujifilm). After fixation, slices were washed for 10 min x 3 times with PBS and then permeabilized with 0.3% Triton X-100 in PBS. Subsequently, slices were treated with 0.3% Triton X-100 and 0.5% BSA in PBS to prevent non-specific staining. Next, they were treated with 0.3% Triton X-100 and streptavidin-Cy3 (1:500) in PBS and were again incubated overnight in 4°C. After washing steps, slices were finally mounted with DAKO S3023 medium, and coverslips were applied immediately. Confocal images were scanned through a 40x water-immersed objective (N.A. 0.5) from FV1200 confocal microscope (Olympus Microscopy). Branch orders were manually counted from a series of z-section images (z step: approx. 1 μm, 512 × 512 pixels) displayed using Fluoview software (FV31S).

### Data availability

All data supporting the results presented in the manuscript were included in the figures.

## Acknowledgments

This study has been supported by National Research Foundation of Korea (2020R1A2C2006438) and Seoul National University Hospital.

## Figure legends

**Fig. S1.**
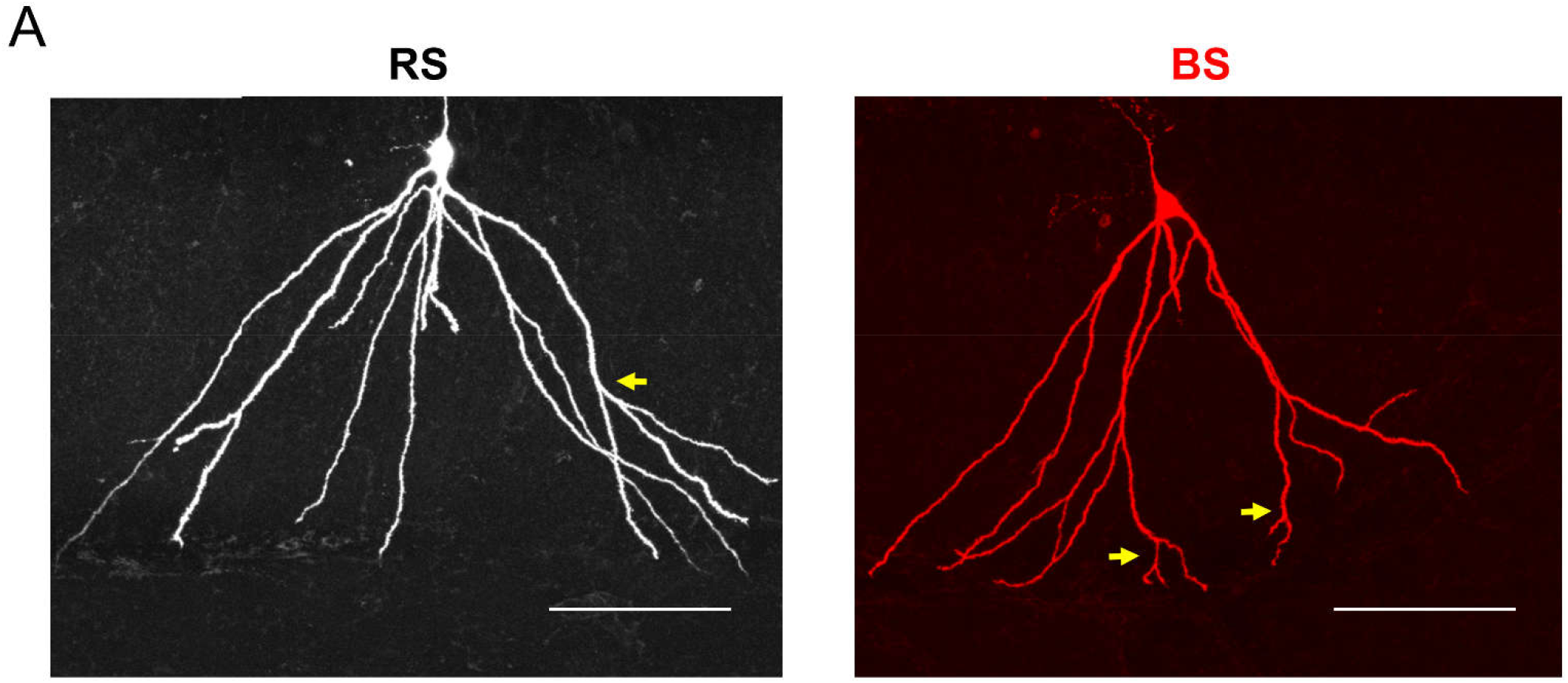
Representative biocytin-filled RS-(*left*) and BS-GC (*right*). Yellow arrows indicate the maximal dendritic branching points. Scale bar, 100 μm.

**Fig. S2.**
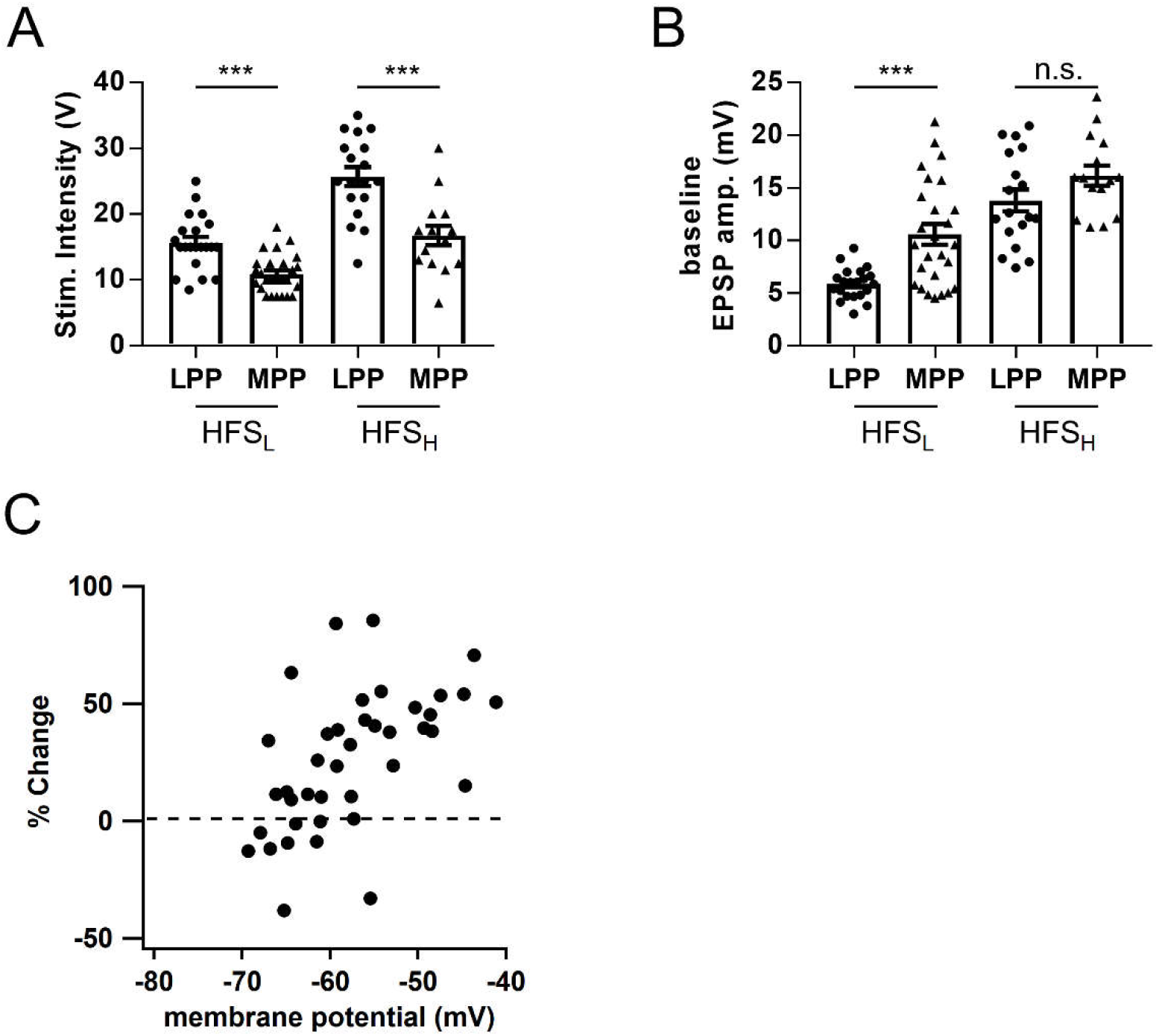
Stimulation Intensities and baseline EPSP amplitudes to evoke sub-or suprathreshold voltage responses at MPP and LPP synapses. **A:** Mean stimulation intensities used for HFS_L_ and HFS_H_ at MPP and LPP synapses. Both mean intensities for HFS_L_ (LPP, 15.6 ± 0.9 V, n = 21; MPP, 10.9 ± 0.6 V, n = 26; ***p<0.001) and HFS_H_ (LPP, 25.7 ± 1.4 V, n = 18; MPP, 16.7 ± 1.5 V, n = 15; ***p<0.001) were significantly stronger at LPP-GCs than MPP-GCs. **B:** Baseline amplitudes of EPSP evoked by HFS_L_ and HFS_H_ at MPP and LPP synapses. Significantly larger baseline EPSP amplitudes were required at MPP-GCs than LPP-GCs in order to elicit subthreshold responses (HFS_L_: LPP, 5.9 ± 0.3 mV, n = 21; MPP, 10.6 ± 1.0 mV, n = 26; ***p<0.001). But, it was not significant to elicit 3 APs responses (HFS_H_: LPP, 13.8 ± 1.0 mV, n = 18; MPP, 16.2 ± 1.0 mV, n = 15; p = 0.18). **C:** Plot of LTP_sub_ magnitude as a function of peak membrane potential of EPSP summation. Error bars, S.E.M. ***p<0.001. n.s., not significant (p>0.05).

**Fig. S3.**
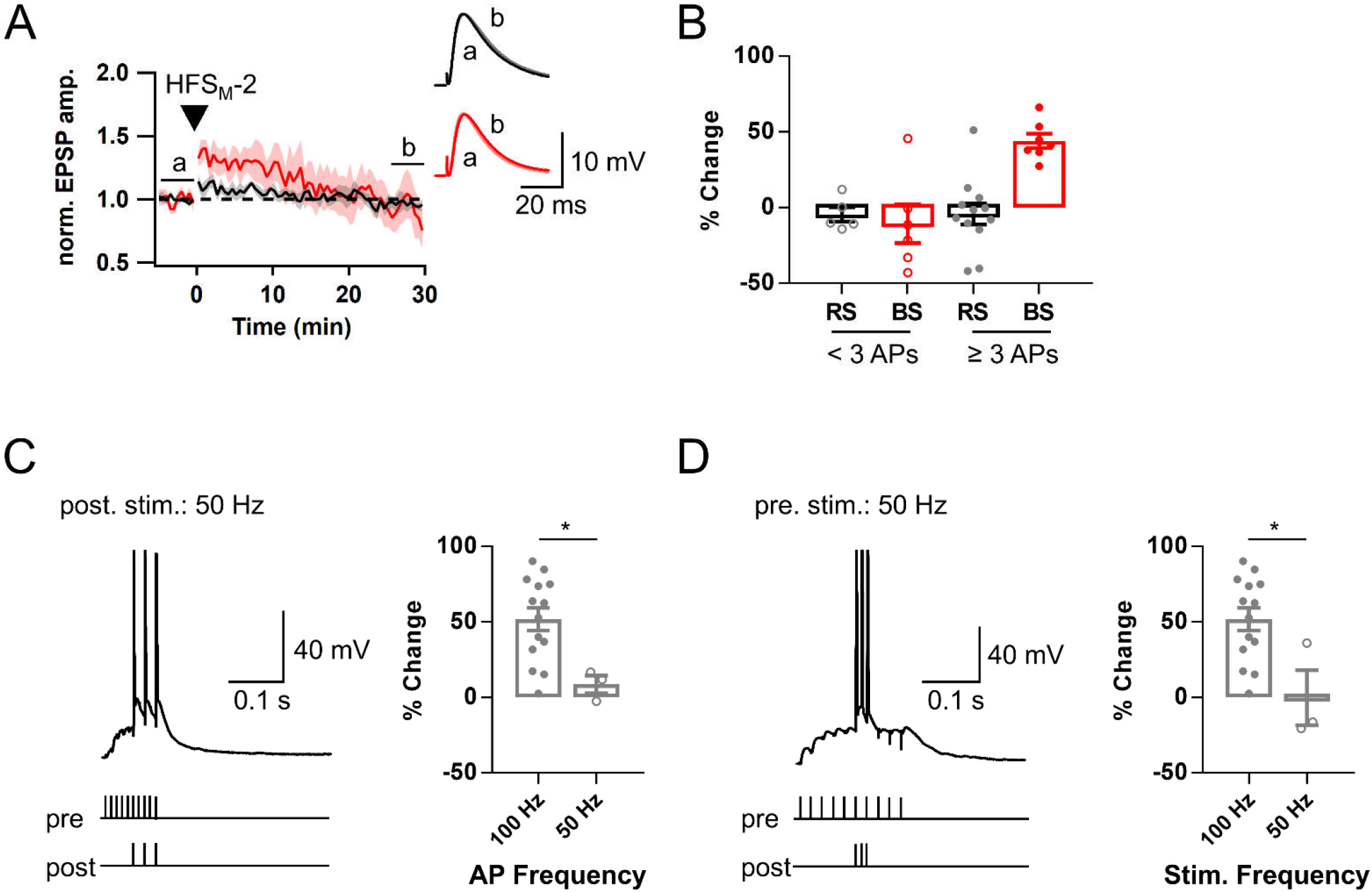
Conditions for LTP_AP_ induction. **A:** Time course of normalized EPSP before and after HFS_M_-2. HFS_M_-2 is defined by HFS eliciting 1 or 2 APs. Note that LTP was not maintained not only in RS (black) but also in BS (red). **B:** Magnitudes of LTP_AP_ induced by HFS_M_ or HFS_H_ in RS and GS (RS, -4.5 ± 4.8%, n = 5; BS, -10.6 ± 12.8 %, n = 6). **C:** Dependence of LTP_AP_ on the postsynaptic AP frequency. When the post-synaptic AP bursts were elicited at 50 Hz instead of 100 Hz in the pairing protocol (*left*), LTP_AP_ was not induced (*right*, 8.7 ± 5.8%, n = 3, *p<0.05). **D:** Dependence of LTP_AP_ on the synaptic stimulation frequency. When ten EPSPs were evoked at 50 Hz instead of 100 Hz in the pairing protocol (*left*), LTP_AP_ was not induced (−0.2 ± 18.2%, n = 3, *p<0.05). Shades and error bars, S.E.M. *p < 0.05.

**Fig. S4.**
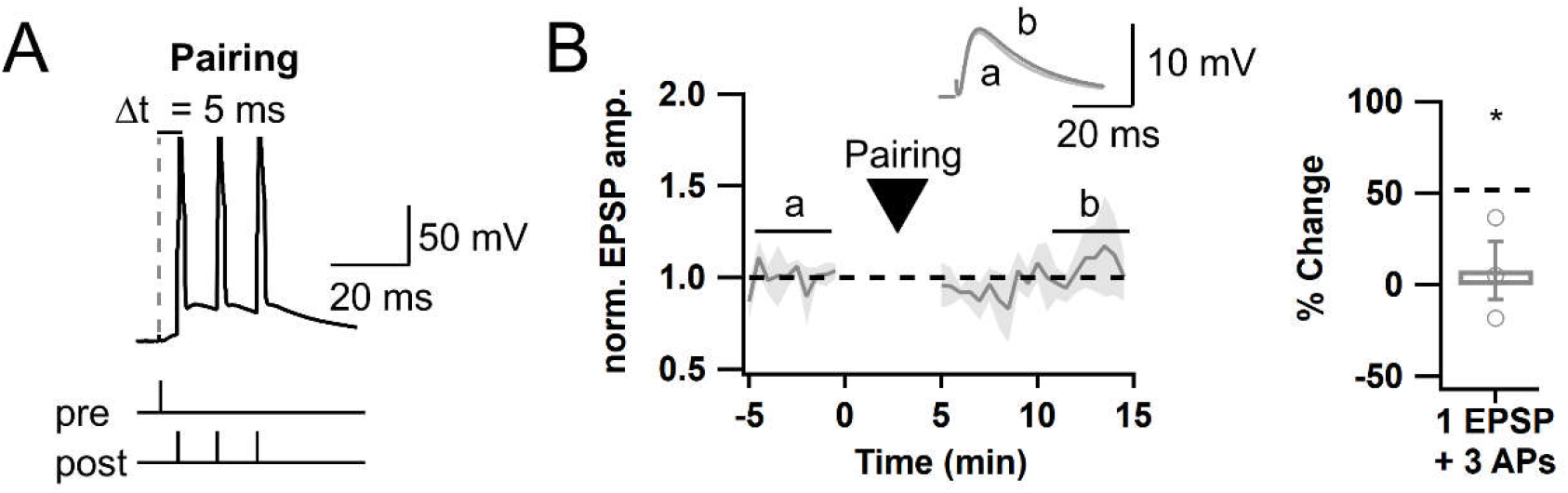
Single presynaptic stimulation is not sufficient to induce LTP. **A:** Representative voltage response to a pairing protocol, in which a single EPSP was coupled to 3 APs at 100 Hz (pre-post time interval = 5 ms). **B:** *Left*, Time course of normalized EPSP before and after applying the pairing protocol shown in *A* 300 times (for 5 min every 1 s). *Right*, LTP_AP_ was not induced by this induction protocol (7.8 ± 15.9%, n = 3, *p<0.05). *Black dashed line*, mean value for LTP magnitude induced by the conventional pairing protocol comprised of 10 EPSPs and 3 APs at 100 Hz as shown in *Figure 3F*. Shades and error bars, S.E.M.

## References

Bayazitov IT, Richardson RJ, Fricke RG & Zakharenko SS. (2007). Slow presynaptic and fast postsynaptic components of compound long-term potentiation. Journal of Neuroscience 27, 11510–11521.

Bittner KC, Milstein AD, Grienberger C, Romani S & Magee JC. (2017). Behavioral time scale synaptic plasticity underlies CA1 place fields. Science 357, 1033–1036.

Chen S, Yue C & Yaari Y. (2005). A transitional period of Ca2+-dependent spike afterdepolarization and bursting in developing rat CA1 pyramidal cells. The Journal of physiology 567, 79–93.

Colino A & Malenka RC. (1993). Mechanisms underlying induction of long-term potentiation in rat medial and lateral perforant paths in vitro. Journal of neurophysiology 69, 1150–1159.

Diamantaki M, Frey M, Berens P, Preston-Ferrer P & Burgalossi A. (2016). Sparse activity of identified dentate granule cells during spatial exploration. Elife 5, e20252.

Dumenieu M, Senkov O, Mironov A, Bourinet E, Kreutz MR, Dityatev A, Heine M, Bikbaev A & Lopez-Rojas J. (2018). The low-threshold calcium channel Cav3. 2 mediates burst firing of mature dentate granule cells. Cerebral Cortex 28, 2594–2609.

Grover LM & Teyler TJ. (1990). Two components of long-term potentiation induced by different patterns of afferent activation. Nature 347, 477–479.

Hsu C-L, Zhao X, Milstein AD & Spruston N. (2018). Persistent sodium current mediates the steep voltage dependence of spatial coding in hippocampal pyramidal neurons. Neuron 99, 147-162. e148.

Hunt DL, Linaro D, Si B, Romani S & Spruston N. (2018). A novel pyramidal cell type promotes sharp-wave synchronization in the hippocampus. Nature neuroscience 21, 985–995.

Kampa BM, Letzkus JJ & Stuart GJ. (2006). Requirement of dendritic calcium spikes for induction of spike-timing-dependent synaptic plasticity. The Journal of physiology 574, 283–290.

Kim S, Jung D & Royer S. (2020). Place cell maps slowly develop via competitive learning and conjunctive coding in the dentate gyrus. Nature communications 11, 1–15.

Kim S, Kim Y, Lee S-H & Ho W-K. (2018). Dendritic spikes in hippocampal granule cells are necessary for long-term potentiation at the perforant path synapse. Elife 7, e35269.

Larkum ME, Kaiser K & Sakmann B. (1999). Calcium electrogenesis in distal apical dendrites of layer 5 pyramidal cells at a critical frequency of back-propagating action potentials. Proceedings of the National Academy of Sciences 96, 14600–14604.

Lee JW & Jung MW. (2017). Separation or binding? Role of the dentate gyrus in hippocampal mnemonic processing. Neuroscience & Biobehavioral Reviews 75, 183–194.

Letzkus JJ, Kampa BM & Stuart GJ. (2006). Learning rules for spike timing-dependent plasticity depend on dendritic synapse location. Journal of Neuroscience 26, 10420–10429.

Lisman J. (2017). Glutamatergic synapses are structurally and biochemically complex because of multiple plasticity processes: long-term potentiation, long-term depression, short-term potentiation and scaling. Philosophical Transactions of the Royal Society B: Biological Sciences 372, 20160260.

Lisman JE. (1997). Bursts as a unit of neural information: making unreliable synapses reliable. Trends in neurosciences 20, 38–43.

Llinás RR & Steriade M. (2006). Bursting of thalamic neurons and states of vigilance. Journal of neurophysiology 95, 3297–3308.

Okubo Y & Iino M. (2011). Visualization of glutamate as a volume transmitter. The Journal of Physiology 589, 481–488.

Pernía-Andrade AJ & Jonas P. (2014). Theta-gamma-modulated synaptic currents in hippocampal granule cells in vivo define a mechanism for network oscillations. Neuron 81, 140–152.

Pignatelli M, Ryan TJ, Roy DS, Lovett C, Smith LM, Muralidhar S & Tonegawa S. (2019). Engram cell excitability state determines the efficacy of memory retrieval. Neuron 101, 274-284. e275.

Raus Balind S, Magó Á, Ahmadi M, Kis N, Varga-Németh Z, Lőrincz A & Makara JK. (2019). Diverse synaptic and dendritic mechanisms of complex spike burst generation in hippocampal CA3 pyramidal cells. Nature communications 10, 1–15.

Remy S & Spruston N. (2007). Dendritic spikes induce single-burst long-term potentiation. Proceedings of the National Academy of Sciences 104, 17192–17197.

Schmidt-Hieber C, Jonas P & Bischofberger J. (2004). Enhanced synaptic plasticity in newly generated granule cells of the adult hippocampus. Nature 429, 184–187.

Schwindt PC & Crill WE. (1995). Amplification of synaptic current by persistent sodium conductance in apical dendrite of neocortical neurons. Journal of neurophysiology 74, 2220–2224.

Wang W, Trieu BH, Palmer LC, Jia Y, Pham DT, Jung K-M, Karsten CA, Merrill CB, Mackie K & Gall CM. (2016). A primary cortical input to hippocampus expresses a pathway-specific and endocannabinoid-dependent form of long-term potentiation. Eneuro 3.

Watanabe S, Hoffman DA, Migliore M & Johnston D. (2002). Dendritic K+ channels contribute to spike-timing dependent long-term potentiation in hippocampal pyramidal neurons. Proceedings of the National Academy of Sciences 99, 8366–8371.

Wu J, Harney S, Rowan MJ & Anwyl R. (2008). Involvement of group I mGluRs in LTP induced by strong high frequency stimulation in the dentate gyrus in vitro. Neuroscience letters 436, 235–238.

Yue C, Remy S, Su H, Beck H & Yaari Y. (2005). Proximal persistent Na+ channels drive spike afterdepolarizations and associated bursting in adult CA1 pyramidal cells. Journal of Neuroscience 25, 9704–9720.

Zhang X, Schlögl A & Jonas P. (2020). Selective routing of spatial information flow from input to output in hippocampal granule cells. Neuron 107, 1212-1225. e1217.

